# ULK1-regulated AMP sensing by AMPK and its application for the treatment of chronic kidney disease

**DOI:** 10.1101/2023.08.09.552390

**Authors:** Tomoki Yanagi, Hiroaki Kikuchi, Koh Takeuchi, Koichiro Susa, Takayasu Mori, Naohiro Takahashi, Takefumi Suzuki, Yuta Nakano, Tamami Fujiki, Yu Hara, Soichiro Suzuki, Yutaro Mori, Fumiaki Ando, Shintaro Mandai, Shinya Honda, Satoru Torii, Shigeomi Shimizu, Tatemitsu Rai, Shinichi Uchida, Eisei Sohara

## Abstract

AMP-activated protein kinase (AMPK) is a central kinase involved in energy homeostasis. Increased intracellular adenosine monophosphate (AMP) levels result in AMPK activation through the binding of AMP to the γ-subunit of AMPK. Recently, we reported that AMP-induced AMPK activation is impaired in the kidneys in chronic kidney disease (CKD) despite an increase in the AMP/ATP ratio. However, the mechanisms by which AMP sensing is disrupted in CKD are unclear. In this study, we identified mechanisms of energy homeostasis in which Unc-51-like kinase 1 (ULK1)-dependent phosphorylation of AMPKγ1 at Ser260/Thr262 promotes AMP sensitivity of AMPK. AMPK activation by AMP was impaired in *Ulk1^−/−^* mice despite an increased AMP/ATP ratio. We also demonstrated that MK8722, an allosteric AMPK activator, activates AMPK in the kidneys of a CKD mouse model via a pathway that is independent of AMP sensing. MK8722 treatment significantly attenuates the deterioration of renal function in CKD and is a potential therapeutic option in CKD therapeutics.

## Introduction

AMP-activated protein kinase (AMPK) is an evolutionarily conserved serine/threonine kinase that is a key player in cellular energy homeostasis (1). When intracellular energy is low (increase in AMP levels), AMPK activates pathways associated with energy (adenosine triphosphate [ATP]) production to maintain energy homeostasis (1,2). AMPK consists of three (α, β, and γ) subunits (3,4) and the α-subunit contains the catalytic domain (5). AMP binding to the γ-subunit causes a conformational change in the β-subunit (6), which facilitates phosphorylation at Thr172 in the α-subunit by AMPK kinases (7), such as LKB1 (8), resulting in AMPK activation. Although AMP binding to the γ-subunit activates AMPK, no studies have identified the physiological regulator of this event other than the AMP/ATP ratio (9–13).

The kidneys are energy-intensive organs that are responsible for electrolyte transport and waste excretion (14). Thus, it is likely that AMPK is involved in energy metabolism in the kidneys (15–18). There is evidence that AMPK is relevant in chronic kidney disease (CKD), a progressive disease that leads to end-stage renal disease and a requirement for renal replacement therapy. AMP sensitivity of AMPK is impaired in kidneys of CKD model mice, resulting in AMPK inactivation (19). Interestingly, AMPK activity was reduced in the CKD model despite an increased AMP/ATP ratio, indicating the presence of an unknown mechanism that regulates AMP sensing by AMPK. AMP insensitivity during CKD may lead to further deterioration of the kidneys, forming a vicious cycle of CKD exacerbation. Elucidation of the detailed molecular mechanisms of failure of AMP sensing is required for an understanding of the mechanisms of CKD progression and the development of novel therapeutic strategies; this is particularly important as therapeutic interventions for CKD are currently limited (20,21).

Unc-51-like kinase 1 (ULK1) is an attractive candidate to explain the reduced AMP-sensing activity of AMPK observed in CKD. Mutual regulation between ULK1 and AMPK has been previously reported (22). ULK1 is activated upon AMPK phosphorylation (23); however, ULK1 also phosphorylates AMPK (24). ULK1 phosphorylation is considerably reduced in the kidneys of mice with CKD (19). Additionally, the suppression of AMPK kinase activity in the kidneys of ULK1 knockout (*Ulk1−/−*) mice resulted in an energy deficit associated with an increased AMP/ATP ratio as well as exacerbated renal dysfunction and fibrosis (25). These suggest that ULK1 deficiency suppresses AMPK activity in the kidneys, and that there is an unknown mechanism through which ULK1 regulates AMP sensing by AMPK.

Herein, we elucidated the molecular mechanisms by which AMPK senses AMP and the mechanisms through which AMP sensitivity is disrupted during CKD and ULK1-knockout conditions. We discovered a mechanism of energy homeostasis in which ULK1 increases the binding of AMP to the AMPK γ-subunit through direct phosphorylation and revealed the mechanisms of AMP-sensing failure by AMPK in the kidneys and in CKD models. Finally, we identified a compound capable of activating AMPK even when AMPK fails to sense AMP. Importantly, treatment with this compound attenuated CKD progression by improving multiple markers of CKD in a mouse model, indicating that it may be a potential therapeutic option for CKD.

## Results

### ULK1 plays a major role in AMP sensing by AMPK

AMPK is activated by phosphorylating Thr172 when cellular energy is low (i.e., when the AMP/ATP ratio is increased). However, AMPKα phosphorylation at Thr172 was downregulated in the kidneys of a 5/6 nephrectomy model (5/6Nx) of CKD mice despite an elevated AMP/ATP ratio (**Figs. 1A, B**; see **Source Data file** for statistics), consistent with our previous study (19). Additionally, both total ULK1 (T-ULK1) and phosphorylated ULK1 (Ser555, P-ULK1) expression were decreased in the kidneys of mice with CKD (**Fig. 1A**). Based on these results, we hypothesized that ULK1 is involved in the impaired AMP sensitivity of AMPK. To test this hypothesis, we analyzed ULK1 knockout (*Ulk1^−/−^*) mice (26) and observed that AMPK was not activated in *Ulk1^−/−^* kidneys during starvation despite an elevated AMP/ATP ratio, similar to the effect observed in CKD kidneys (**Figs. 1C, D**; see **Source Data file** for statistics). These results suggest that AMPK does not correctly sense AMP under ULK1-deficient conditions.

**Fig. 1.**
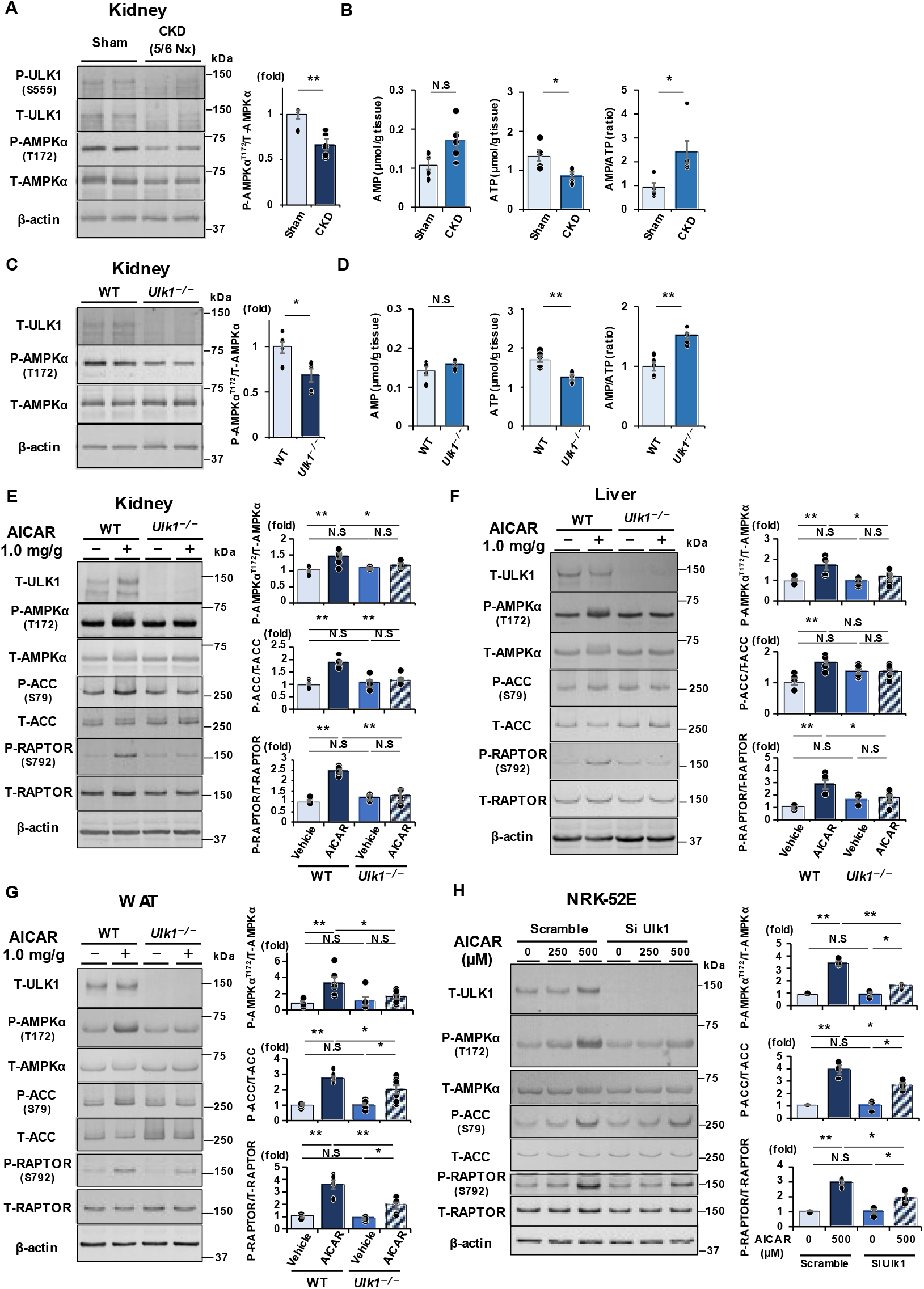
Unc-51-like kinase 1 (ULK1) plays a major role in AMP sensing by AMP- activated protein kinase (AMPK) **A.** Representative immunoblots of total and phosphorylated ULK1, and total AMPKα and phospho-AMPKα^Thr172^ expression, in the kidneys of sham and CKD (5/6 nephrectomy) mice (n = 5). **B.** AMP/ATP ratio in the kidneys of WT mice in the sham and CKD groups (n = 5). **C.** Representative immunoblots of total AMPKα and phospho-AMPKα^Thr172^ expression in kidneys of wild-type (WT) or *Ulk1^−/−^* mice following starvation (n = 5). **D.** AMP/ATP ratio in kidneys of WT or *Ulk1^−/−^* mice (n = 5). **E.** Representative immunoblots and densitometric analysis of P-AMPKα^Thr172^, P- ACC^Ser79^, and P-RAPTOR^Ser792^ in kidneys of WT or *Ulk1^−/−^*mice treated intraperitoneally with or without 5-aminoimidazole 4-carboxamide-1-beta-d- ribofuranoside (AICAR) (1.0 mg/g BW) (n = 6). **F.** Representative immunoblots and densitometric analysis of P-AMPKα^Thr172^, P-ACC^Ser79^, and P-RAPTOR^Ser792^ expression in liver tissue of WT or *Ulk1^−/−^*mice treated with or without AICAR (n = 6). **G.** Representative immunoblots and densitometric analysis of P-AMPKα^Thr172^, P- ACC^Ser79^, and P-RAPTOR^Ser792^ in white adipose tissue of WT or *Ulk1^−/−^* mice treated with or without AICAR (n = 6). **H.** Representative immunoblots and densitometric analysis of P-AMPKα^Thr172^, P- ACC^Ser79^, and P-RAPTOR^Ser792^ expression in NRK-52E cells transfected with control siRNA (scramble) or ULK1 siRNA (Si-Ulk1) with or without AICAR for 12 h (n = 3). *P* values (\**P* < 0.05, \*\**P* < 0.01) were calculated using the two-tailed unpaired *t*-test or one-way analysis of variance with Tukey’s honestly significant difference test. Data are presented as the mean ± standard error of the mean. For detailed data, statistical analysis, and exact *P* values, refer to the **Source Data file**.

To confirm our hypothesis, 5-aminoimidazole 4-carboxamide-1-beta-D-ribofuranoside (AICAR), an AMP analog, was injected intraperitoneally (1.0 mg/g body weight) into both wild-type (WT) and *Ulk1^−/−^* mice, and changes in AMPK activation (upregulated AMPKα phosphorylation at Thr172) in the kidneys were evaluated. Upregulation of AMPKα^Thr172^ phosphorylation induced by AICAR was observed in WT mice but not in *Ulk1^−/−^* mice (**Fig. 1E**; see **Source Data file** for statistics). This indicates that the AMP-sensing mechanism is dysregulated in *Ulk1^−/−^* mice, suggesting that ULK1 plays a major role in the AMP-sensing mechanism of AMPK. Phosphorylation of acetyl-CoA carboxylase (ACC) at Ser 79, which is a downstream protein of AMPK that is involved in fatty acid oxidation, and phosphorylation of Raptor at Ser792, which is directly phosphorylated by activated AMPK (27), were not upregulated in *Ulk1^−/−^*mice (**Fig. 1E**). AICAR-induced AMPK activation was also absent in both liver (**Fig. 1F**; see **Source Data file** for statistics) and white adipose tissue (WAT) (**Fig. 1G**; see **Source Data file** for statistics) of *Ulk1*^-/-^ mice. These findings indicate that AICAR-induced dysregulation of AMPK activation in *Ulk1^−/−^* mice is not restricted to the kidneys and reveals a broad role of ULK1 in AMP-dependent energy metabolism in multiple organ systems.

To confirm the direct effects of ULK1 on AMP sensing by AMPK, we performed RNA interference experiments in cultured renal tubular epithelial (NRK-52E) cells. We confirmed that the absence of ULK1 impairs AMP sensitivity of AMPK in NRK-52E renal tubular epithelial cells (**Fig. 1H**; see **Source Data file** for statistics). AICAR-induced upregulation of AMPKα^Thr172^ phosphorylation was attenuated by ULK1 depletion in NRK-52E cells. These results indicate that ULK1 deficiency results in impaired AMP-induced AMPK activation both in vivo and in vitro, and that ULK1 plays an important role in the AMP-sensing mechanism of AMPK, which is dysregulated during CKD.

### ULK1 binds to AMPKγ1 and directly phosphorylates it at Ser260 and Thr262

Although ULK1 deficiency impaired AMP sensing by AMPK, the underlying mechanisms remain unknown. Reportedly, AMPKγ directly binds to AMP through CBS domains (6,28,29), enabling a conformational change in the β-subunit (6) and leading to α- subunit phosphorylation at Thr172 and subsequent AMPK activation (7). Therefore, we focused on the effect of ULK1 on the AMPK γ-subunit. As expected, co-immunoprecipitation experiments revealed a direct interaction between AMPKγ1 and ULK1 (**Fig. 2A**). Previously, mass spectrometry revealed that Ser260, Thr262, and Ser269 of AMPKγ1 were phosphorylated by ULK1 (22); however, the functional significance of phosphorylation at these sites was not reported. As shown in **Fig. 2B**, Ser260 and Thr262 of AMPKγ1 are evolutionarily conserved in mammals. To confirm whether these amino acids are phosphorylated by ULK1, we generated an anti-phospho- AMPKγ1 (Ser260/Thr262) antibody (**Fig. S1)** and performed kinase assays. As shown in **Fig. 2C** and **Fig. S2,** recombinant ULK1 and trimeric AMPKα1β1γ1 resulted in the direct phosphorylation of AMPKγ1 at Ser260 and Thr262 by ULK1. AMPKγ1 phosphorylation was not confirmed using the mutant trimeric protein AMPKα1β1γ1^S260A/T262A^ (**Fig. 2C**). AMPKγ1 phosphorylation was also observed when a purified AMPKγ1 monomer and ULK1 were co-incubated, demonstrating that ULK1 directly phosphorylates AMPKγ1 at Ser260 and Thr262 (**Fig. 2D**). Furthermore, AMPKγ1^Ser260/Thr262^ phosphorylation was upregulated in AMPKγ1 bound to ULK1 (**Fig. 2E**). These results suggest that ULK1 binds to and directly phosphorylates AMPKγ1 at Ser260 and Thr262.

**Fig. 2.**
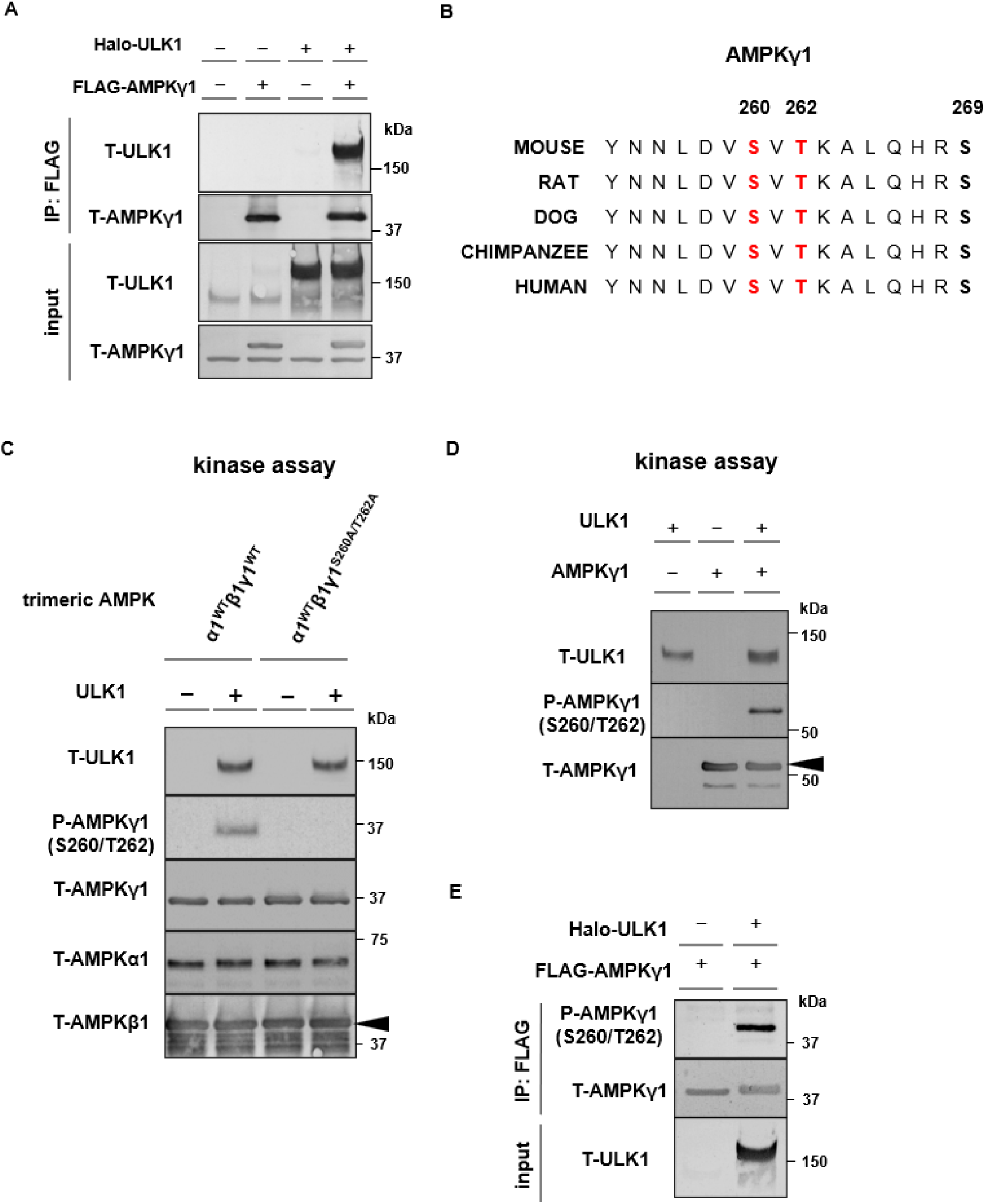
ULK1 binds with AMPKγ1 and directly phosphorylates at Ser260/Thr262. **A.** Immunoprecipitation of FLAG-tagged AMPKγ1 and/or Halo-tagged ULK1 expression in HEK293T cells showing direct interaction between AMPKγ1 and ULK1. **B.** Evolutionally conserved sequence alignment of AMPKγ1 in mammals. **C.** Immunoblot of proteins subjected to a kinase assay using purified trimeric AMPK consisting of AMPKγ1 wild-type (WT) or AMPKγ1^S260A/T262A^ mutant with and without ULK1 showing direct phosphorylation of AMPKγ1 at S260/T262 by ULK1. **D.** Immunoblots of proteins subjected to a kinase assay using purified monomeric AMPKγ1 and ULK1. **E.** Immunoprecipitation of FLAG-tagged AMPKγ1 and/or Halo-tagged ULK1 expression in HEK293T cells showing AMPKγ1 bound to T-ULK1 is likely to be phosphorylated at S260/T262.

### ULK1 promotes AMPKα activation through AMPKγ1 phosphorylation at Ser260/Thr262

We determined whether phosphorylated AMPKγ1^Ser260/Thr262^ promotes AMPK kinase activity. AMPK catalytic activity was measured according to the phosphorylation level of AMPKα at Thr172, a critical site for AMPK activation (30). ULK1 overexpression upregulated AMPKγ1^Ser260/Thr262^ phosphorylation, and phosphorylated AMPKγ1 was bound more readily to phosphorylated AMPKα at Thr172 compared with nonphosphorylated AMPKα (**Fig. 3A**; see **Source Data file** for statistics). Additionally, the AMPKγ1^S260A/T262A^ mutant exhibited decreased binding to phosphorylated AMPKα (**Fig. 3B**; see **Source Data file** for statistics). To confirm the effects of ULK1 on AMPK catalytic activity, we examined the effects of ULK1 activation on AMPK. We used the selective ULK1 agonist BL918 (31) to human embryonic kidney 293 (HEK293) cells and found that BL918 upregulated ULK1 phosphorylation at Ser555 and enhanced AMPKγ1^Ser260/Thr262^ phosphorylation (**Fig. 3C**; see **Source Data file** for statistics). Similar to the findings in **Fig. 3A**, AMPKγ1, which was phosphorylated in the presence of BL918, was more likely to bind to phosphorylated AMPKα^Thr172^ (**Fig. 3D**; see **Source Data file** for statistics). Furthermore, this activating effect of BL918 on AMPKα (AMPKα phosphorylation at Thr172) was diminished in the AMPKγ1^S260A/T262A^ mutant (**Fig. 3D**; see **Source Data file** for statistics) and in ULK1-knockdown-cultured NRK- 52E cells (**Fig. 3E**; see **Source Data file** for statistics), indicating that ULK1 facilitates AMPKα activation through AMPKγ1 phosphorylation at Ser260/Thr262.

**Fig. 3.**
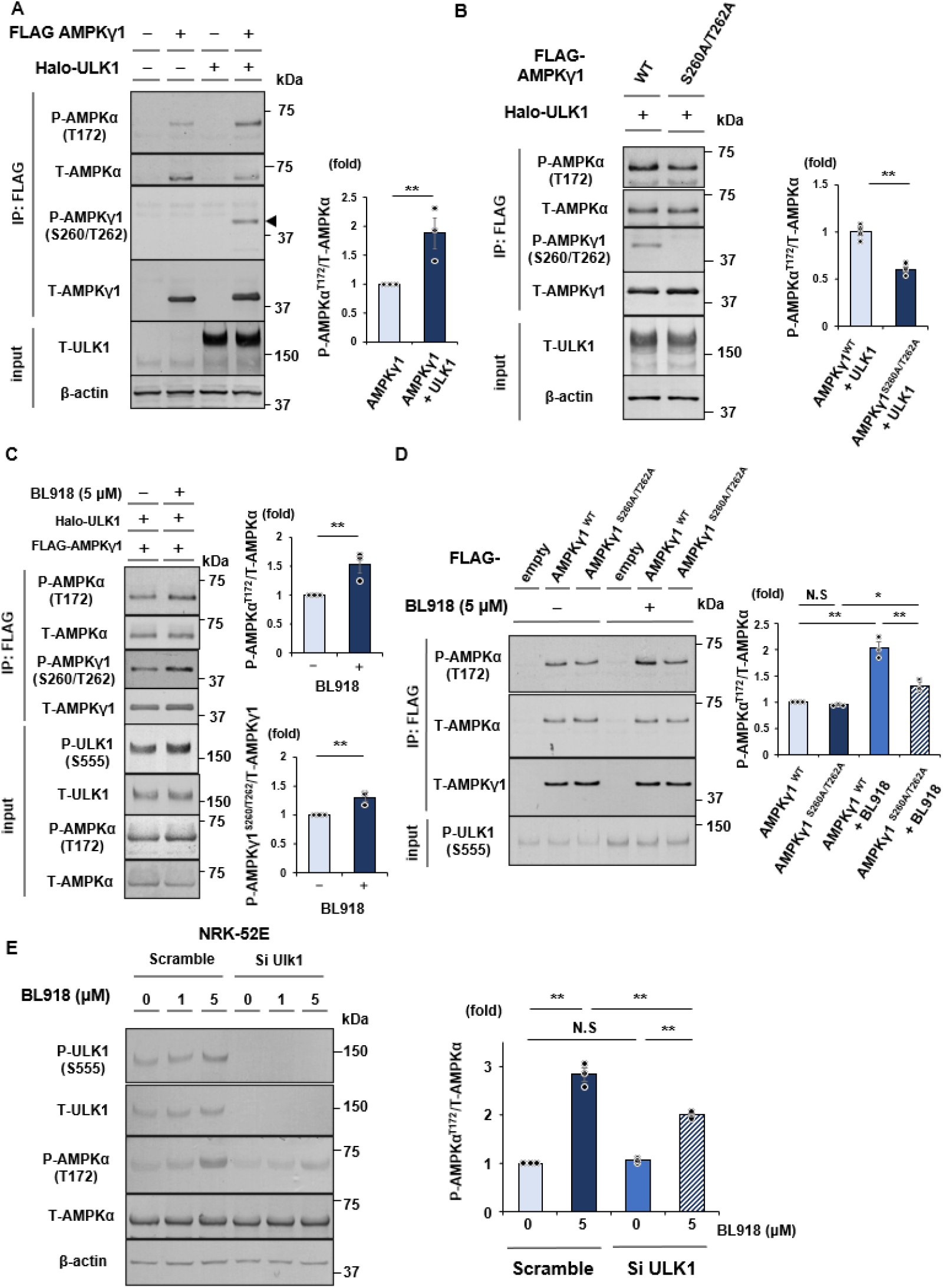
Overexpression or activation of ULK1 activates the AMP-activated protein kinase alpha subunit (AMPKα) through AMPKγ1^Ser260/Thr262^ phosphorylation. **A.** Immunoprecipitation of FLAG-tagged AMPKγ1 with/without Halo-tagged ULK1 expression in HEK293T cells showing AMPK bound to T-ULK1 is likely to be phosphorylated at P-AMPKα^Thr172^. Representative immunoblots and densitometric analysis of P-AMPKα^Thr172^ expression when ULK1 is overexpressed (n = 3). **A. B.** Immunoprecipitation of FLAG-tagged AMPKγ1^WT^ or AMPKγ1^S260A/T262A^ with Halo- tagged ULK1 expression in HEK293T cells. Representative immunoblots and densitometric analysis of P-AMPKα^Thr172^ complexed with AMPKγ1^WT^ or AMPKγ1^S260A/T262A^ when ULK1 is overexpressed (n = 4). **B.** Immunoprecipitation of FLAG-tagged AMPKγ1 with Halo-tagged ULK1 expression in HEK293T cells treated with/without BL918 (selective ULK1 activator) for 15 minutes. Representative immunoblots and densitometric analysis of P-AMPKγ1^Ser260/Thr262^ (lower) and P-AMPKα^Thr172^ (upper) expression (n = 3). **C.** Immunoblots and densitometric analysis of P-AMPKα^Thr172^ complexed with AMPKγ1^WT^ or AMPKγ1^S260A/T262A^ with/without 15 minutes BL918 (selective ULK1 activator, 5 μM) treatment (n = 3). **D.** Representative immunoblots and densitometric analysis of P-AMPKα^Thr172^ expression in NRK-52E cells transfected with control siRNA (scramble) or ULK1 siRNA (Si-Ulk1) with/without BL918 for 15 minutes (n = 3). *P* values (**\****P* < 0.05, **\*\****P* < 0.01) were calculated using the two-tailed unpaired *t*-test or one-way analysis of variance with Tukey’s honestly significant difference test. Data are presented as the mean ± standard error of the mean. For detailed data, statistical analysis, and exact P values, refer to the **Source Data file**.

### ULK1-knockout and CKD model mice exhibited downregulated phosphorylation of AMPKγ1 at Ser260/Thr262 and AMPKα at Thr172 *in vivo*

We confirmed AMPKγ1^Ser260/Thr262^ phosphorylation by ULK1 *in vivo* using tissues from WT and *Ulk1^−/−^* mice. Mice were starved to avoid the effects of dietary regulation on AMPK. AMPKγ1^Ser260/Thr262^ and AMPKα^Thr172^ phosphorylation was downregulated in kidney and liver tissues of starved *Ulk1^−/−^* mice compared with those in WT mice (**Figs. 4A, B**; see **Source Data file** for statistics). We subsequently evaluated whether AMPKγ1^Ser260/Thr262^ phosphorylation was downregulated in the kidneys of CKD mice. Expectedly, AMPKγ1^Ser260/Thr262^ phosphorylation was also downregulated in the kidneys of 5/6Nx model CKD mice relative to control mice (**Fig. 4C**; see **Source Data file** for statistics).

**Fig. 4.**
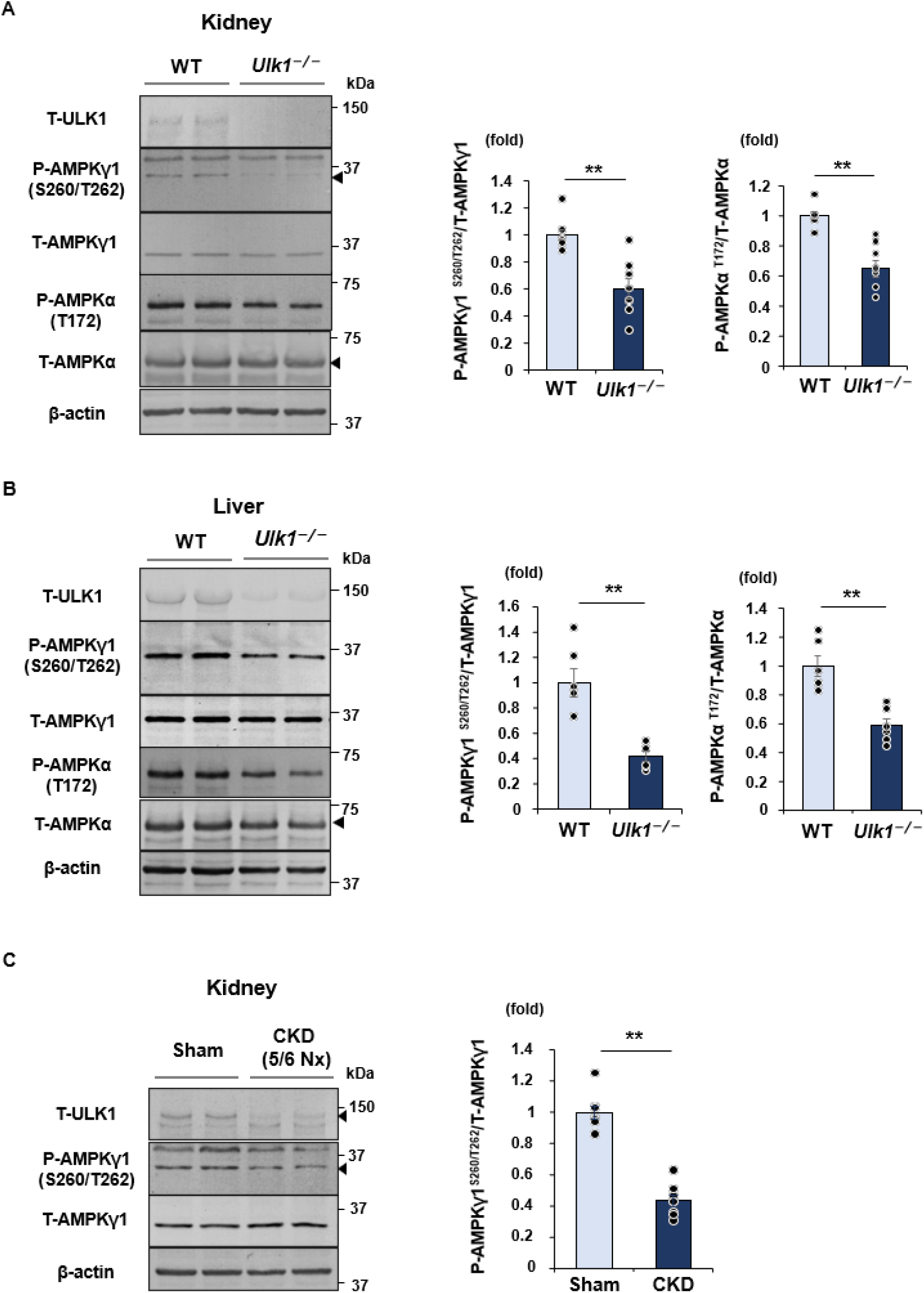
ULK1-knockout mice and CKD model mice exhibit decreased phosphorylation of AMP-activated protein kinase gamma-1 (AMPKγ1) at Ser260/Thr262 and of AMPKα at Thr172. **A.** Representative immunoblots and densitometric analysis of P-AMPKγ1^Ser260/Thr262^ and P-AMPKα^Thr172^ expression in kidneys of wild-type (WT) or *Ulk1^−/−^* mice following starvation (n = 8). **B.** Representative immunoblots and densitometric analysis of P-AMPKγ1^Ser260/Thr262^ and P-AMPKα^Thr172^ expression in livers of WT or *Ulk1^−/−^* mice following starvation (n = 6). **C.** Representative immunoblots and densitometric analysis of P-AMPKγ1^Ser260/Thr262^ expression in kidneys of mice subjected to sham operation or 5/6 nephrectomy to induce chronic kidney disease (CKD) (n = 8). *P* values (**\****P* < 0.05, **\*\****P* < 0.01) were calculated using a two-tailed unpaired *t*-test. Data are presented as the mean ± standard error of the mean. For detailed data, statistical analysis, and exact P values, refer to the **Source Data file**.

**Fig. 5.**
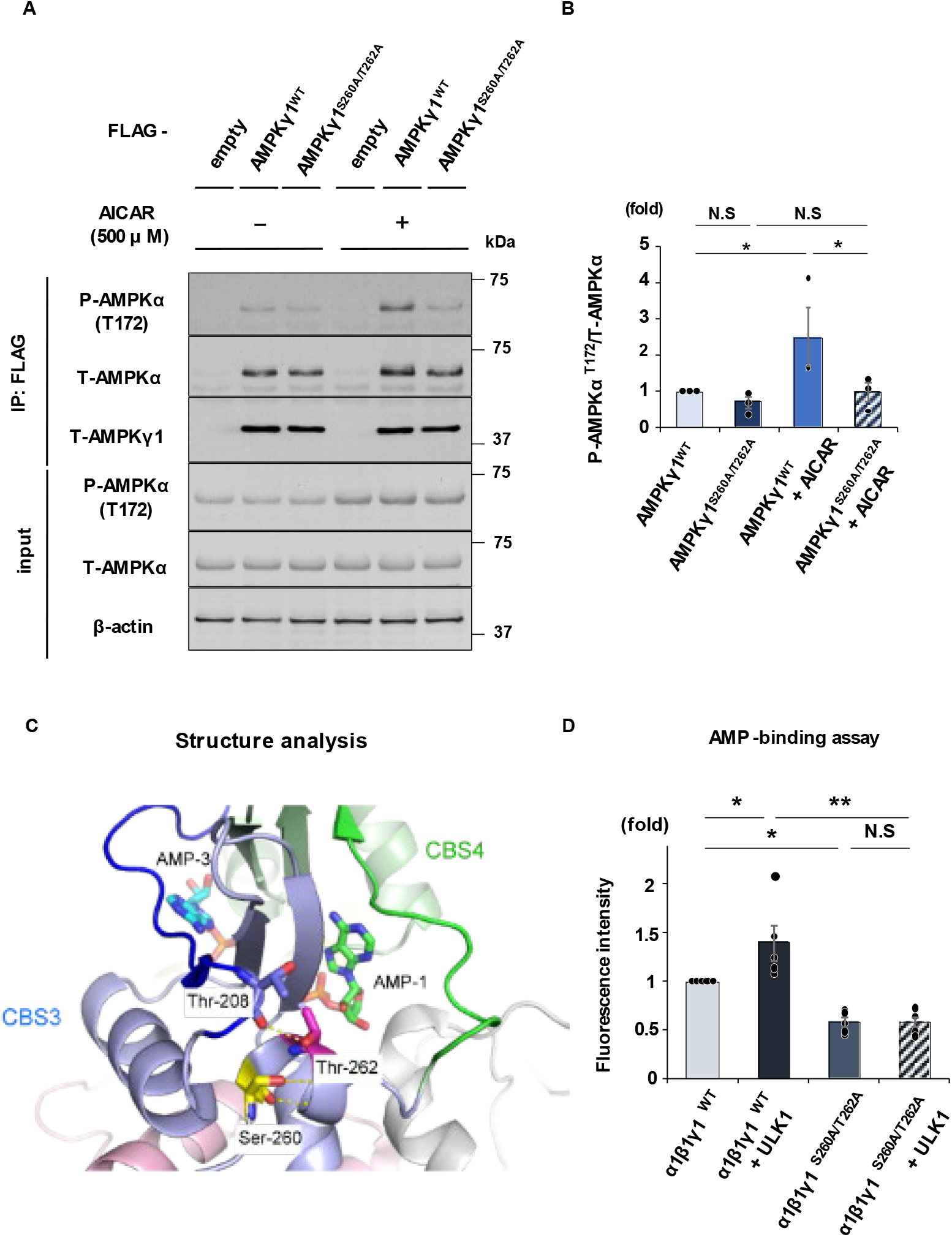
AMPKγ1^Ser260/Thr262^ phosphorylation increases the affinity of AMPKγ1 for AMP. **A, B.** Representative immunoblots (A) and densitometric analysis (B) evaluating the sensitivity of P-AMPKα^Thr172^ complexed with AMPKγ1 wild-type (WT) or S260A/T262A mutant to 12h-AICAR treatment (n = 3). **A. C.** Crystal structure revealing the role of Ser260 and Thr262 in maintaining the structure of AMPKγ1. **B. D.** Fluorescence intensity of Mant-AMP (fluorescent dye) bound to purified trimeric AMPK protein consisting of AMPKγ1^WT^ or AMPKγ1^S260A/T262A^ with/without ULK1 (n = 5) *P* values (**\****P* < 0.05) were calculated using one-way analysis of variance with Tukey’s honestly significant difference test. Data are presented as the mean ± standard error of the mean. For detailed data, statistical analysis, and exact P values, refer to the **Source Data file**.

### AMPKγ1^Ser260/Thr262^ phosphorylation by ULK1 increases the affinity of AMPKγ1 for AMP

Next, we determined whether AMPKγ1^Ser260/Thr262^ phosphorylation by ULK1 regulates AMP sensing of AMPK. We evaluated AMPKα phosphorylation at Thr172 in WT AMPKγ1 and phospho-deficient S260A/T262A mutants in the presence of AICAR in a cultured cell line. AMPKα co-immunoprecipitated by phospho-deficient AMPKγ1^S260A/T262A^, did not exhibit upregulated Thr172 phosphorylation with AICAR treatment. In contrast, AMPKα co-immunoprecipitated by WT AMPKγ1 showed upregulated phosphorylation at Thr172 with AICAR treatment, indicating that the AMPKγ1^S260A/T262A^ mutant is less sensitive to increased AMP levels (**Figs. 5A, B**; see **Source Data file** for statistics). For comparison, AMPKα bound to AMPKγ1^S269A^, which was mutated at a different ULK1 phosphorylation site, did not exhibit a decreased response relative to controls upon AICAR treatment, similar to the findings for AMPKα bound to WT AMPKγ1 (**Fig. S3)**. This finding suggests that AMPKγ1 phosphorylation at Ser269 is not involved in AMP sensing by AMPK. The results also indicate that AMPKγ1 phosphorylation at Ser260/Thr262 is directly involved in the AMP-sensing mechanism of AMPK, and that phosphorylation at these sites by ULK1 markedly affects AMPK kinase activity in response to AMP.

To investigate the significance of AMPKγ1 phosphorylation at Ser260/Thr262, we examined the structure of the AMPK γ-subunit. Although there is no structural information available for dual-phosphorylated AMPKγ, the phosphorylation sites are in a structurally important loop connecting the CBS3 and CBS4 domains in the γ-subunit (PDB ID 4CFH, **Fig. 5C**). In the crystal structure we generated, Ser260 Oγ and Thr262 Oγ formed hydrogen bonds with Lys263 HN and Thr208 CO, respectively, and stabilized the conformation of the adenine nucleotide-binding loops in the CBS3 and CBS4 domains (shown in blue and green, respectively), thereby forming the so-called AMP-1 and AMP-3 sites, respectively (32). Therefore, dual phosphorylation of Ser260 and Thr262 most likely affects the conformation of the AMP-binding loops and modifies the affinity of AMPK for AMP as well as enhances its AMP-sensing activity.

From this structural analysis, we hypothesized that AMPKγ^Ser260/Thr262^ phosphorylation directly regulates the amount of AMP binding to the CBS domains. To determine whether these phosphorylation sites are involved in AMP binding to the γ- subunit, we generated trimeric AMPKα1^WT^β1^WT^γ1^WT^ (AMPKα1β1γ1^WT^) and AMPKα1^WT^β1^WT^γ1^S260A/T262A^ mutant (AMPKα1β1γ1^S260A/T262A^) protein complexes (**Fig. 2C**) and co-incubated them with the ULK1 protein to evaluate the binding of AMP to the AMPK constructs by labeling them with a fluorescent dye (Mant-AMP). Recombinant AMPKγ1 phosphorylation at Ser260 and Thr262 by ULK1 significantly increased the binding of AMPK to AMP compared with that in controls (**Fig. 5D**; see also **Supplementary Table 1, Source Data file** for statistics). In contrast, AMP binding to the AMPKα1β1γ1^S260A/T262A^ construct was significantly lower compared with that of AMPKα1β1γ1^WT^, revealing that the AMPK S260A/T262A mutation reduces AMP binding, which suggests that the phosphorylation state of AMPK at these sites is critical for this process. Furthermore, the effects of ULK1, which increases the binding of AMP to AMPKγ1, were not observed in AMPKα1β1γ1^S260A/T262A^ (**Fig. 5D**; see also **Supplementary Table 1, Source Data file** for statistics), suggesting that increased binding of AMP to AMPKγ1 induced by ULK1 is mediated by AMPKγ1 phosphorylation at Ser260 and Thr262. Thus, ULK1 regulates the affinity of AMPKγ1 for AMP through phosphorylation at Ser260/Thr262.

### MK8722 activates AMPK signals in ULK1 deficiency

Our data indicate that decreased ULK1-dependent sensing of AMP by the AMPK γ-subunit contributes to reduced AMP sensing in CKD. Therefore, AMPK activation independent of ULK1 regulation (or AMPK activation that bypasses the AMP-sensing mechanism) may improve CKD by enhancing energy homeostasis in the kidneys. We identified MK8722, and allosteric AMPK activator, as a candidate for such a treatment. MK8722 is a small-molecule compound that directly activates AMPK by binding to an allosteric drug and metabolite site, which is distinct from the sites that bind to AMP, located in a pocket between the AMPKα and AMPKβ subunits. Thus, MK8722 can activate AMPK in a manner that bypasses the AMP-sensing mechanism (33,34). MK8722- activated AMPKα co-immunoprecipitated phospho-deficient AMPKγ1^S260A/T262A^ to comparable levels as those of WT AMPKγ1 (**Fig. 6A**; see **Source Data file** for statistics). This result is in contrast to that following AICAR treatment wherein the mechanism is by AMP binding (**Fig. 5A**). Additionally, MK8722 activates AMPK in NRK-52E cells following ULK1 depletion (**Fig. 6B**; see **Source Data file** for statistics). Furthermore, we determined the in vivo effect of MK8722 and observed upregulated AMPKα phosphorylation at Thr172 in the kidneys of *Ulk1^−/−^* mice treated with MK8722 (**Fig. 6C**; see **Source Data file** for statistics). These results demonstrate that MK8722 activates AMPK even under ULK1-deficient conditions.

**Fig. 6.**
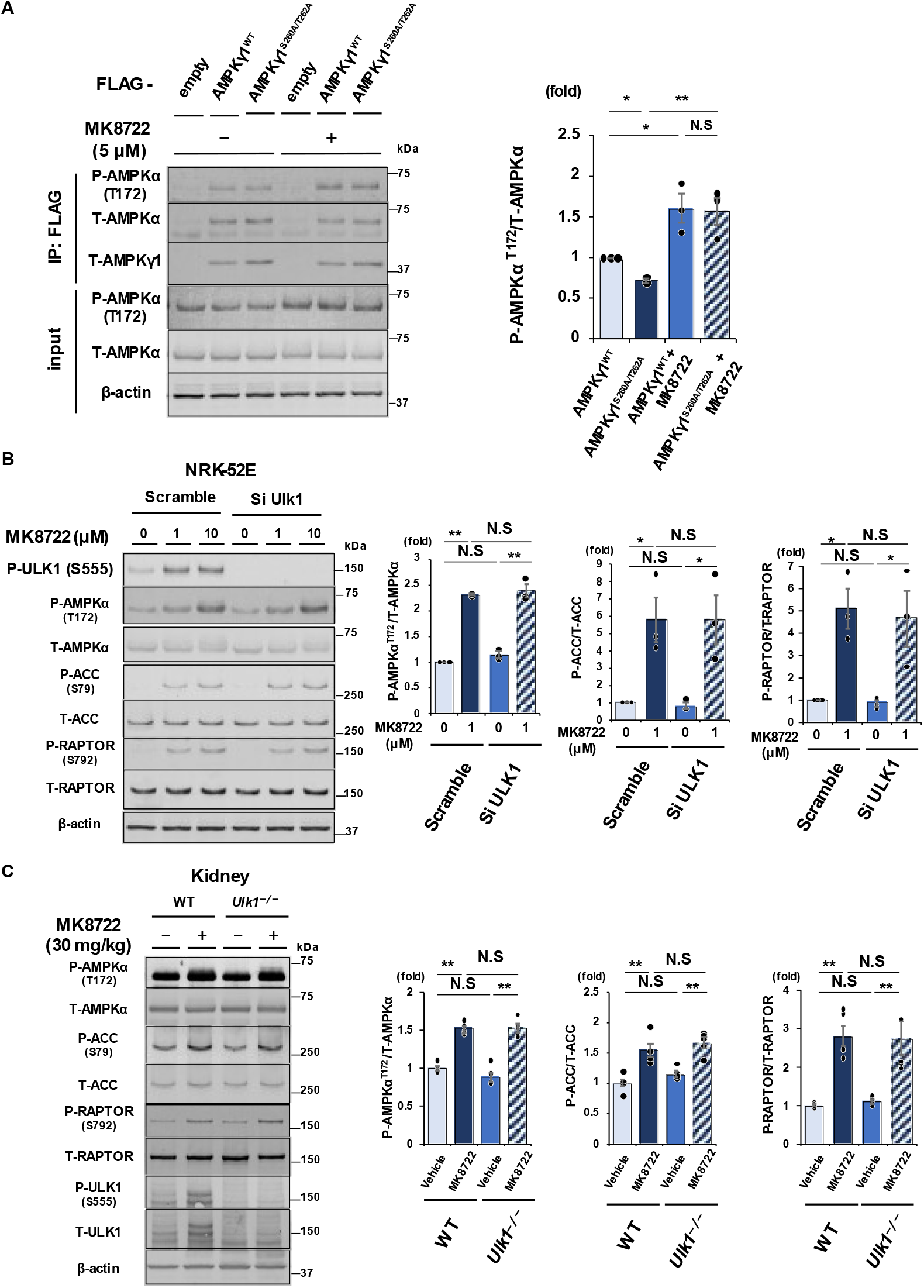
MK8722 directly activates AMP-activated protein kinase alpha (AMPKα) at Thr172 even in the absence of ULK1. **A.** Representative immunoblots and densitometric analysis of P-AMPKα^Thr172^ complexed with AMPKγ1 wild-type (WT) or AMPKγ1^S260A/T262A^ treated with/without MK8722 (n = 3). **B.** Representative immunoblots and densitometric analysis of P-AMPKα^Thr172^, P- ACC^Ser79^, and P-RAPTOR^Ser792^ expression in NRK-52E cells transfected with control siRNA (scramble) or ULK1 siRNA (Si-Ulk1) with or without MK8722 treatment (n = 3). **C.** Representative immunoblots and densitometric analysis of P-AMPKα^Thr172^, P- ACC^Ser79^, and P-RAPTOR^Ser792^ expression in kidneys of WT or *Ulk1^−/−^* mice treated intraperitoneally with or without MK8722 at 30 mg/kg BW (n = 5). *P* values (**\****P* < 0.05, **\*\****P* < 0.01) were calculated using the two-tailed unpaired *t*-test or a one-way analysis of variance with Tukey’s honestly significant difference test. Data are presented as the mean ± standard error of the mean. For detailed data, statistical analysis, and exact P values, refer to the **Source Data file**.

### MK8722 attenuates CKD progression

To determine the effect of MK8722 on CKD progression, MK8722 was administered to CKD mice with a 5/6 nephrectomy model. Treatment with MK8722 (10 mg/kg BW i.p. daily) for 14 days resulted in significantly increased AMPK kinase activity and upregulated phosphorylation of ACC, which is a downstream signal of AMPK in the kidneys (**Fig. 7A**; see **Source Data file** for statistics). Additionally, the levels of renal fibrosis factors collagen types 1 alpha (*Col1a*) and 3 alpha (*Col3a*) and actin alpha-2 (*Acta2*) significantly decreased in MK8722-treated CKD mice (**Fig. 7B**). CKD mice treated with MK8722 showed significantly lower serum creatinine levels and preserved renal function (**Fig. 7C**; see **Source Data file** for statistics). Histologically, kidneys treated with MK8722 exhibited reduced renal fibrosis in a 5/6 nephrectomy model (**Fig. 7D**). These results indicate that MK8722 attenuates CKD progression through AMPK activation.

**Fig. 7.**
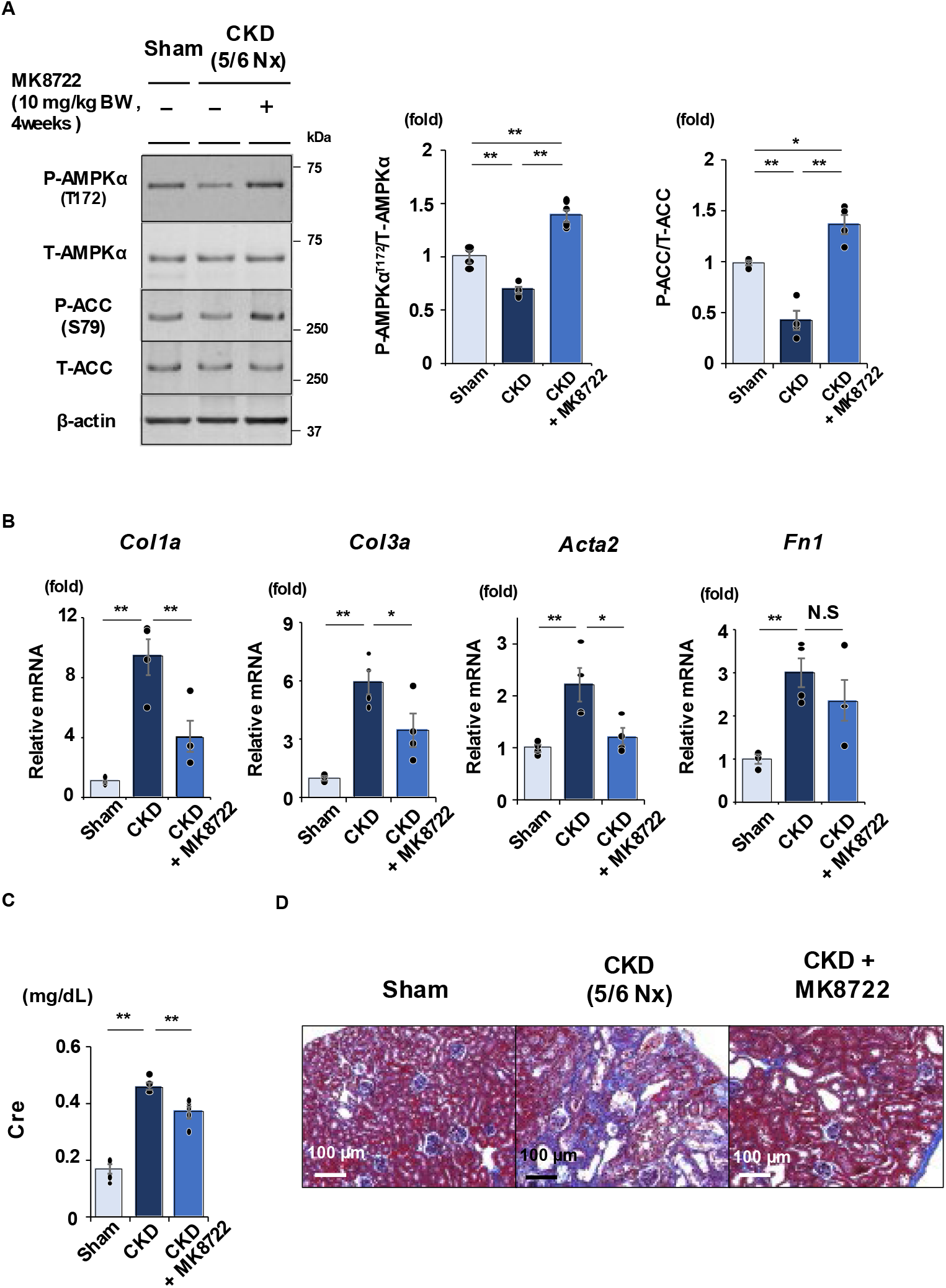
MK8722 attenuates the progression of chronic kidney disease (CKD) **A.** Representative immunoblots and densitometric analysis of P-AMPKα^Thr172^ and P- ACC^Ser79^ expression in mouse kidneys after sham operation, 5/6 nephrectomy (to induce CKD), or CKD treated with MK8722 (10 mg/kg BW) (n = 4). **B.** Quantitative reverse transcription-polymerase chain reaction of pro-fibrotic markers in kidneys of sham, CKD, and CKD treated with MK8722 mice (n = 4). *Col1a*, collagen type 1 alpha; *Col3a*, collagen type 3 alpha; *Acta2*, actin alpha-2; *Fn1*, fibronectin 1. **C.** Serum creatinine (Cre) value of Sham, CKD, and CKD treated with MK8722 mice (n = 4) at 4 weeks after 5/6 nephrectomy. **D.** Masson trichrome staining of mouse kidneys after sham operation, CKD (5/6 nephrectomy), and CKD treated with MK8722. *P* values (**\****P* < 0.05, **\*\****P* < 0.01) were calculated using one-way analysis of variance with Tukey’s honestly significant difference test. Data are presented as the mean ± standard error of the mean. For detailed data, statistical analysis, and exact P values, refer to the **Source Data file**.

**Fig. 8.**
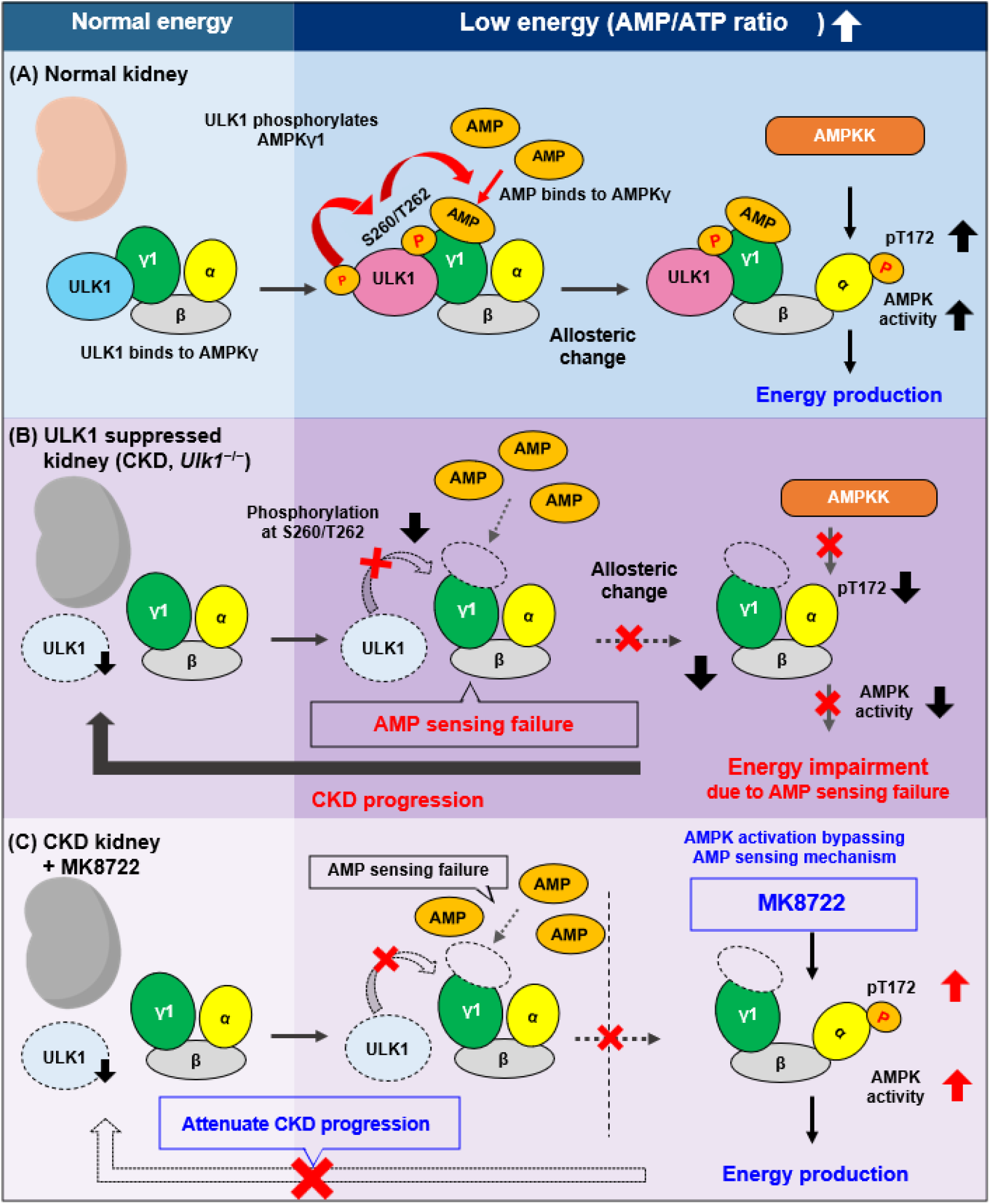
Mechanism by which ULK1 regulates AMP sensing by AMP-activated protein kinase (AMPK) Schematic representation of our findings. **A.** In the presence of ULK1, ULK1 increases AMP sensing of AMPKγ1 through phosphorylation at Ser260/Thr262 in a low-energy state. Consequently, AMPK is subjected to an allosteric conformational change, and AMPKα is highly phosphorylated at Thr172 by AMPK kinase (LKB1). **B.** In the absence of ULK1, phosphorylation of AMPKγ1 at Ser260/Thr262 is impaired, resulting in reduced AMP sensing. Consequently, AMPKα phosphorylation is diminished. Without ULK1, cells cannot efficiently sense AMP and activate AMPK, resulting in a vicious cycle of chronic kidney disease (CKD). **C.** MK8722 activates AMPK bypassing this ULK1-mediated AMP-sensing mechanism, promoting energy production and attenuating CKD progression.

## Discussion

In CKD, damaged nephrons undergo atrophy, while the remaining intact nephrons partially compensate for the loss in function, resulting in increased single-nephron GFR and increasing energy demand by the individual proximal tubule cells. Activation of AMPK, which responds to low cellular energy states, could be a potential treatment for CKD wherein cellular energy is low. Sensing of the AMP/ATP ratio by AMPK is reduced in CKD (19), but the underlying mechanisms have not been elucidated. In this study, we revealed that ULK1 promotes cellular AMP sensing by AMPK through phosphorylation of Ser260/T262 on the AMPKγ1 subunit. Furthermore, phosphorylation of these sites promoted phosphorylation of the α1 subunit of AMPK at Thr172, thereby enhancing AMPK activity. Finally, we demonstrated that the allosteric pan-AMPK activator MK8722 activated AMPK in a manner that bypassed the ULK1-mediated AMP-sensing mechanism. Treatment with MK8722 improved markers for CKD in a mouse model, thus presenting potential candidate for the treatment of CKD (**Figs. 6**, **7**).

BL918, which is a ULK1 activator, may also be a potential therapeutic option for CKD through the improvement of AMP sensitivity of AMPK. Although BL918 activated AMPK in NRK-52E cells (35), ULK1 activation in the kidneys *in vivo* was not confirmed because of the low distribution of BL918 (**Fig. S4 A**). However, AMPK was activated by the ULK1 activator in *ex-vivo* kidney and liver tissues (**Figs. S4 B** and **C)**. These results suggest that ULK1 activation could be a potential treatment option for diseases wherein AMPK is inhibited, including liver diseases wherein AMPK activation is suppressed such as nonalcoholic steatohepatitis (NASH) (36). Future studies are needed to scrutinize the effects of more efficient ULK1 activators with higher bioavailability than BL918.

The binding of AMP to the CBS domains of AMPKγ is dependent on the intracellular energy state (AMP/ATP ratio). Using the direct AMP-binding assay, we confirmed that the ULK1-induced upregulation in binding between AMP and AMPKγ1 is mediated by AMPKγ1 phosphorylation at Ser260 and Thr262 (**Fig. 5).** These results suggest that AMPKγ phosphorylation is important for the structural conformation of the CBS domain, increasing the AMP-binding capacity of AMPKγ. Although AMP binding to the AMPKα1β1γ1^S260A/T262A^ construct was significantly lower than that in WT constructs, AMP binding was not completely deleted by this mutation, suggesting that AMP-sensing mechanisms in AMPK is not restricted to the mechanism we discovered in this study. Future studies are needed to determine how the structure of the CBS loop is altered by phosphorylation.

AMP-sensing failure was observed not only in the kidneys but also in other organs such as the liver and WAT. Thus, AMP sensing by AMPK might be relevant to diseases of other tissues aside from the kidneys. For example, AMPK is highly expressed in adipose tissue (37); cardiac and skeletal muscles (38); liver (39); and immune cells, including T cells (40). Meanwhile, AMPK is inactivated in specific liver diseases and diabetes mellitus (36,37). Furthermore, AMPK is involved in autoimmune disease such as rheumatoid arthritis (RA), in which T cells have an altered metabolic profile wherein glucose is shifted from glycolysis and ATP production to the pentose phosphate pathway and biosynthesis (40). Thus, the potential of AMPK-activating therapies to treat several diseases, such as diabetes mellitus, metabolic liver diseases (such as NASH), or RA, warrants further studies.

Although ULK1 is involved in autophagy (41), the effects of its absence on canonical autophagy in mice vary in a tissue-dependent manner. For example, ULK1 deficiency affects autophagy in mouse thymus tissue (42,43), weakly affects the erythroid lineage (26,42), and has no effect on skeletal muscles (44). Canonical autophagy is reportedly maintained in the kidneys of ULK1 knockout mice (25). We also demonstrated that there are no significant changes in LC3-I and LC3-II levels or SQSTM/p62 expression in the kidneys of *Ulk1^−/−^*and WT mice (25). These results indicate that canonical autophagy is not involved in the renal phenotype of *Ulk1^−/−^*mice and that the regulatory mechanism of AMPK through ULK1 is independent of autophagy.

This study demonstrated that ULK1 phosphorylates AMPK and significantly affects AMPKα kinase activity by using cultured cells and a mouse model. In contrast, activated ULK1 inhibited AMPKα phosphorylation at Thr172 via negative feedback (22,24). Although the exact reason for this discrepancy is not entirely clear, organ or cell type specificity may account for the differences in the effects of ULK1. However, we detected reduced AMP sensing in several tissues from ULK1-knockout mice, suggesting that ULK1 promotes the response of AMPK to increased AMP levels. Further studies should clarify whether this role of ULK1 varies among tissue types.

We focused on AMPKγ1 in this study despite the AMPKγ subunit having three isoforms (γ1, γ2, and γ3). AMPKγ1 and AMPKγ2 are expressed in all organs, whereas AMPKγ3 is specifically expressed in skeletal muscles (45). As AMPKγ1 is a major regulatory subunit of AMPK activity (11), we focused on the γ1 subunit. However, considering that AMPKγ2, but not AMPKγ3, also has conserved phosphorylation sites (Thr490/Thr492) similar to Ser260/Thr262 of AMPKγ1 (**Fig. S5**), AMPKγ2 could also be involved in the AMP-sensing mechanism of AMPK.

In summary, we identified mechanisms by which ULK1 regulates AMP sensitivity. ULK1 regulates AMP sensitivity of AMPK by phosphorylating AMPKγ, thereby regulating the affinity of AMPKγ for AMP. This finding provides evidence of the physiological regulation of AMP sensitivity by AMPKγ. We also demonstrated that MK8722 allosterically activates AMPK to bypass the AMP-sensing mechanism. These findings may lead to novel treatment strategies for CKD.

## Methods

### Cell culture

HEK293 cells and NRK-52E rat kidney epithelial cells were purchased from the American Type Culture Collection (Manassas, VA, USA); both cell lines were cultured in Dulbecco’s Modified Eagle’s Medium containing 4.5 g/L glucose (Nacalai Tesque, Tokyo, Japan) supplemented with 5% or 10% fetal bovine serum at 37°C and 5% CO2 in a humidified incubator. AICAR (Fujifilm Wako Pure Chemical Industries, Osaka, Japan) was used at concentrations of 250 and 500 μM according to a previous study (19). Cell samples were collected 12 h after administering AICAR. MK8722 (MedChemExpress, Monmouth Junction, NJ, USA) was used at concentrations of 1, 5, and 10 μM according to a previous study (34), and samples were collected 15 min after loading. BL918 (Selleck Chemicals, Houston, TX, USA) was used at concentrations of 1 and 5 μM according to a previous study (31), and cell samples were collected 15 minutes after loading.

#### Plasmid and siRNA

Polymerase chain reaction (PCR) was used to isolate mouse AMPKγ1 cDNA from a mouse AMPKγ1-HA vector (#40605, Addgene, Watertown, MA, USA). Mouse AMPKγ1 cDNA was then cloned into the 3xFLAG-CMV10 vector (Sigma-Aldrich, St. Louis, MO, USA). Mouse ULK1 cDNA was isolated from a p3xFLAG-CMV14-mULK1 vector (#24301, Addgene) by PCR and cloned into the Halo-tagged pFN21A vector (Promega, Madison, WI, USA). The AMPKγ1 mutant vector wherein Ser260 and Thr262 were replaced with alanine was created using parent-template backbone mouse AMPKγ1- full-length vector (# 15996, Addgene) and primers with the desired mutation. The parent template is removed using a methylation-dependent endonuclease. For protein expression and purification experiments, mouse cDNA was isolated from AMPKγ1 and S260A/T262A-mutant AMPKγ1 by PCR using the aforementioned AMPKγ1, pAMPK- alpha1-full-length (#27297, Addgene), and AMPK-β1-FLAG (#40602, Addgene) vectors. Each mouse cDNA was then cloned into a pGEX-6p-1 vector (Cytiva, Marlborough, MA, USA) with the sequence of each construct being confirmed. For ULK1 depletion experiments, Ulk1 Rat siRNA Oligo Duplex (SR514693, Origene, San Francisco, CA, USA) was used. According to the manufacturer’s instructions, Lipofectamine™ RNAiMAX kit (Invitrogen) was used to transfect 10 nM si-ULK1 into HK-2 cells for experiments. For the accuracy of the transfection, negative control siRNA was transfected into HK-2 cells in si-NC group for 48 h. After completing the above operation, we incubated the cells in the DMEM/F12 medium containing 5% FBS for 36h. Western-blot analysis detected the level of target protein in the si-ULK1 group to verify the effect of siRNA transfection.

### Protein expression and purification

Recombinant GST fusion mouse AMPKγ1 (WT or S260A/T262A mutant), AMPKα1 (WT), and AMPKβ1 were expressed in *Escherichia coli* BL21 cells and purified using glutathione-sepharose beads (46). GST fusion proteins were eluted by cleaving the GST tag with a precision protease (Cytiva, Marlborough, MA, USA).

### Animal experiments

All C57BL/6 mice used in this study were purchased from CLEA Japan (Tokyo, Japan). Meanwhile, all *Ulk1^−/−^* mice in this study were obtained from Dr. Shimizu in the Department of Pathological Cell Biology, Tokyo Medical and Dental University (Tokyo, Japan) (26). All animal experiments were approved by the Animal Care and Use Committee of Tokyo Medical and Dental University and conducted in accordance with the animal experiment guidelines of the Japanese Ministry of Education, Culture, Sports, Science and Technology. Mice aged 6–8 weeks underwent a sham operation or 5/6 Nx (CKD model mice) (19); 4 weeks after, blood samples were collected from the venous plexus near the mandible and analyzed using iSTAT EC8 (Abbott, Inc., Abbott Park, IL, USA) and iSTAT CHEM8 (Abbott, Inc.). For the AICAR loading experiment, mice were intraperitoneally administered with AICAR (Fujifilm Wako Pure Chemical Industries, Osaka, Japan) at 1.0 mg/g body weight; 45 min after intraperitoneal injection, the mice were sacrificed. For the MK8722 loading experiment, mice were intraperitoneally administered with MK8722 (MedChemExpress) at 30 mg/kg body weight and sacrificed 30 min after injection. To evaluate the therapeutic effects of MK8722 on CKD, CKD mice were administered with MK8722 at 10 mg/kg/day for 4 weeks after 5/6 Nx. To evaluate the effects of BL918 on the kidney, mice were intraperitoneally administered with BL918 at 20 or 40 mg/kg/day and sacrificed 30 mins after injection. In other in vivo experiments, mice were fasted for 12–24 h to eliminate the effects of food intake.

### Ex-vivo kidney and liver tissues experiment

Kidneys and livers were harvested and cut into <500-μm slices using a microslicer (Natsume Seisakusho Co., Ltd., Tokyo, Japan). The sliced samples were then incubated for equivalation in ice-cold Hank’s buffer medium (pH 7.4, 110 mM NaCl, 3 mM KCl, 1.2 mM MgSO4, 1.8 mM CaCl2, 4 mM Na acetate, 1 mM Na citrate, 6 mM d-glucose, 6 mM L-alanine, 1 mM NaH2PO4, 3 mM Na2HPO4, 25 mM NaHCO3) for 20 min at room temperature. Slices from the same kidney or liver were separated and incubated in medium with or without BL918 (5 μM) for 30 min at 28 °C. All solutions were continuously aerated with 95% O2 and 5% CO2. After incubation, the slices were snap frozen in liquid nitrogen and prepared for immunoblotting.

### Immunoblotting

Protein samples were obtained from mouse kidneys, livers, WAT, and cells were cultured for immunoblotting (19). The relative intensities of the immunoblot bands were quantitatively determined using ImageJ software (National Institutes of Health, Bethesda, MD, USA). Primary antibodies used included anti-β-actin (1:1,000, AAN01, Cytoskeleton, Inc., Denver, CO, USA), anti-AMPKα (1:1,000, #2532, Cell Signaling Technology, Danvers, MA, USA), anti-phospho-AMPKα (pThr172) (1:1,000, #2535, Cell Signaling Technology), anti-AMPKβ1 (1:1,000, ab32112, Abcam, Cambridge, UK), anti-AMPKγ1 (1:1,000, ab32508, Abcam), anti-ULK1 (1:1,000, #8054, Cell Signaling Technology), anti-phospho-ULK1 (pSer555) (1:1,000, #5869, Cell Signaling Technology), anti-ACC (1:1,000, #3676, Cell Signaling Technology), anti-phospho-ACC (pSer79) (1:1,000, #3661, Cell Signaling Technology), anti-Raptor (1:1000, #2280, Cell Signaling Technology), anti-phospho-Raptor (Ser792) (1:1,000, #2083, Cell Signaling Technology), and anti-phospho-AMPKγ1. Secondary antibodies used included alkaline phosphatase-conjugated anti-rabbit IgG antibody (1:1,000, #S3738, Promega) and anti-mouse IgG antibody (1:1,000, #S3721, Promega).

### Quantitative reverse transcription (qRT)-PCR

Total RNA obtained from mouse kidneys was reverse-transcribed using ReverTra Ace (TOYOBO, Osaka, Japan). qRT-PCR was performed using the Thermal Cycler Dice Real Time System (Takara Bio, Kusatsu, Japan). Primers and templates were mixed with TB Green Premix Ex Taq II (Takara Bio). mRNA levels were then normalized to that of *Gapdh* and calculated using the comparative Ct method. Sequences for qRT-PCR primers are listed in **Supplementary Table 2**.

### Generation of anti-phospho-AMPKγ1^Ser260/Thr262^ antibody

To create a phospho-specific antibody against AMPKγ1^Ser260/Thr262^, phosphorylated peptide C+NNLDV(pS)V(pT)KALQ was used to immunize rabbits. The antibody produced was obtained and purified from antiserum using phosphorylated peptide-conjugated SulfoLink-coupling resin and dialyzed against PBS. The antibody was further purified by passage through nonphosphopeptide-conjugated SulfoLink- coupling resin. The peptides and antibody were prepared by Eurofins Genomics (Tokyo, Japan).

### In vitro kinase assay using recombinant protein

In total, 1,000 ng of purified AMPKα1 (WT), AMPKβ1 (WT), or AMPKγ1 (WT or S260A/T262A mutation) protein and/or 500 ng of ULK1 protein (MyBioSource, #MBS515701) were prepared and placed in a MOPS-based reaction buffer (25 mM MOPS, pH 7.0, 12.5 mM β-glycerophosphate, 25 mM MgCl2, 5 mM EGTA, 2 mM EDTA, 0.25 mM DTT). The mixture was then incubated at 30°C for 30 min with 200 μM AMP and 50 μM ATP. Samples were denatured for 20 min at 60°C in SDS sample buffer (Cosmo Bio) and subjected to immunoblotting.

### In vitro kinase assay using immunoprecipitated protein from cell lines

3xFLAG-AMPKγ1 and Halo-ULK1 vectors were transfected into HEK293 cells using Lipofectamine 2000. The cells were then collected by immunoprecipitation with anti-FLAG^®^ M2 Affinity Gel (Sigma, A2220) 48 h after transfection. The samples were denatured for 20 min at 60°C with SDS sample buffer and subjected to immunoblotting.

### Measurement of AMP levels in mouse kidneys

AMP levels in kidney tissues were measured using an AMP assay kit (Abcam). Kidneys were harvested from mice and rapidly frozen in liquid nitrogen. After determining the appropriate weight, the tissues were excised and homogenized in AMP buffer. Samples were then centrifuged at 10,000 × *g* and 4°C for 10 min, and the supernatant was collected and incubated with the Reaction Mix at 37°C for 60 min. AMP levels were then calculated by measuring the absorbance at 570 nm using a FilterMax F5 (Molecular Devices, San Jose, CA, USA). A calibration curve with dilutions of AMP was prepared.

### Measurement of ATP levels in mouse kidneys

ATP levels in kidney tissues were measured using the AMERIC-ATP(T) kit (Americ). Kidneys were harvested from mice and rapidly frozen in liquid nitrogen. After determining the appropriate weight, the tissues were excised and homogenized with a phenol-containing reagent. A chloroform-containing reagent was added, and the supernatant was collected after centrifugation at 10,000 × *g* and 4°C for 5 min. The supernatant was diluted, and 10 µL of the sample solution was mixed with 90 µL of luciferase reagent. The maximum bioluminescence intensity was measured using FilterMax F5. A calibration curve with dilutions of ATP was prepared. Data are expressed as ATP μmol/g of tissue.

### AMP-binding assay

Recombinant GST-AMPKγ1 protein was expressed in *Escherichia coli* BL21 cells. Then, 20-mL cultures were grown at 37°C in 2YT broth (1.6% [w/v] tryptone/1% [w/v] yeast extract/0.5% NaCl) containing 100 μg/mL ampicillin until the absorbance at 600 nm was 0.6. Isopropyl-β-D-1-thiogalactopyranoside (1 mM) was added, and cells were cultured for an additional 16 h at 28°C. Cells were isolated by centrifugation and resuspended in 800 μL of ice-cold MOPS buffer followed by sonication (UD-201, Tomy Seiko, Tokyo, Japan). Then, 1% Triton + 0.5% sarkosyl was added to each sample. Samples were incubated at 4°C for 60 min and centrifuged at 4°C for 5 min at 10,000 × *g*. The supernatant was treated with the Slide-A-Lyzer™ Dialysis Cassette Kit (#66382, Thermo Fisher Scientific, Waltham, MA, USA), added to 400 µL of glutathione-sepharose beads (Cytiva), incubated at 4°C for 60 min, recentrifuged, and finally discarded. The remaining glutathione-sepharose beads were incubated at 30°C for 60 min in MOPS buffer containing 1,000 ng of purified AMPKα1 or AMPKβ1 protein, 200 µM Mant-AMP (#NU236, Jena Bioscience, Jena, Germany), 50 µM ATP, and/or 3,000 ng of ULK1 protein (#MBS515701, MyBioSource, San Diego, CA, USA). The beads were then added to Pierce™ centrifuge columns (#89868, Thermo Fisher Scientific) and washed several times with MOPS buffer. The GST tag from purified GST-AMPKγ was cleaved using a cleavage buffer (50 mM Tris-HCl, 150 mM NaCl, 1 mM EDTA, 1 mM DTT) containing precision protease (Cytiva) and eluted. The amount of bound AMP was measured by the binding amount of AMPKγ eluted with Mant-AMP. The fluorescence intensity of Mant-AMP was analyzed using FLUOstar OPTIMA FL (BMG LABTECH, Ortenberg, Germany), with excitation at 355 nm and emission at 460 nm. All experiments were replicated five times.

### Structural analysis

A structural figure was generated using the PyMOL program (ver. 1.8.2.0 Open-source).

### Statistics and reproducibility

Statistical significance was evaluated using the two-tailed unpaired *t*-test. For multiple comparisons, one-way analysis of variance with Tukey’s honestly significant difference test was performed. *P < 0.05 and **P < 0.01 were considered statistically significant. Data are presented as mean ± standard error of the mean. All statistical analyses were performed using JMP^®^ 15 software (SAS Institute Inc., Cary, NC, USA). Source data are provided as a **Source Data file**.

### Study approval

All animal experiments were approved by the Animal Care and Use Committee of Tokyo Medical and Dental University and conducted in accordance with the animal experiment guidelines of the Japanese Ministry of Education, Culture, Sports, Science and Technology (A2023-109A).

#### Author contributions

T.Y. and H.K. contributed equally to this work, and either has the right to list himself first in bibliographic documents. H.K., T.Y., S.U., and E.S. designed the research.

T.Y. and H.K performed experiments. T.Y., H.K., N.T., K.S., T.M., T.F., Y.H, S.S., Y.M., F.A., S.M., T.R., S.U., and E.S. participated in the discussions and interpretation of the data. S.H., S.T., and S.S. generated *Ulk1^−/−^* mice. K.T. performed structural analysis. H.K. and T.Y. wrote the manuscript with input from E.S. and all other authors.

## Acknowledgments

We are grateful to Ms. Motoko Chiga and Ms. Chieko Iijima for their technical assistance. Additionally, the authors thank Yolanda L. Jones; Brigit S. Sullivan, MLS; NIH Library; and Benjamin Carter, Ph. D for editing assistance. This work was supported by Grant Number 21ek0109554h0001 to E. Sohara Japan Agency for Medical Research and Development (AMED), Grant Number 19ek0109304h0002 to S. Uchida Japan Agency for Medical Research and Development (AMED), Grant-in-Aid for Scientific Research (B; 19H03672) and Grant-in-Aid for Challenging Exploratory Research (22K19518) to E. Sohara, Grant-in-Aid for Scientific Research (A; 19H01049) to S. Uchida, Japan Kidney Association-Nippon Boehringer Ingelheim Joint Research Program to E. Sohara, and the Salt Science Research Foundation (2030 and 2131) to E. Sohara.

---

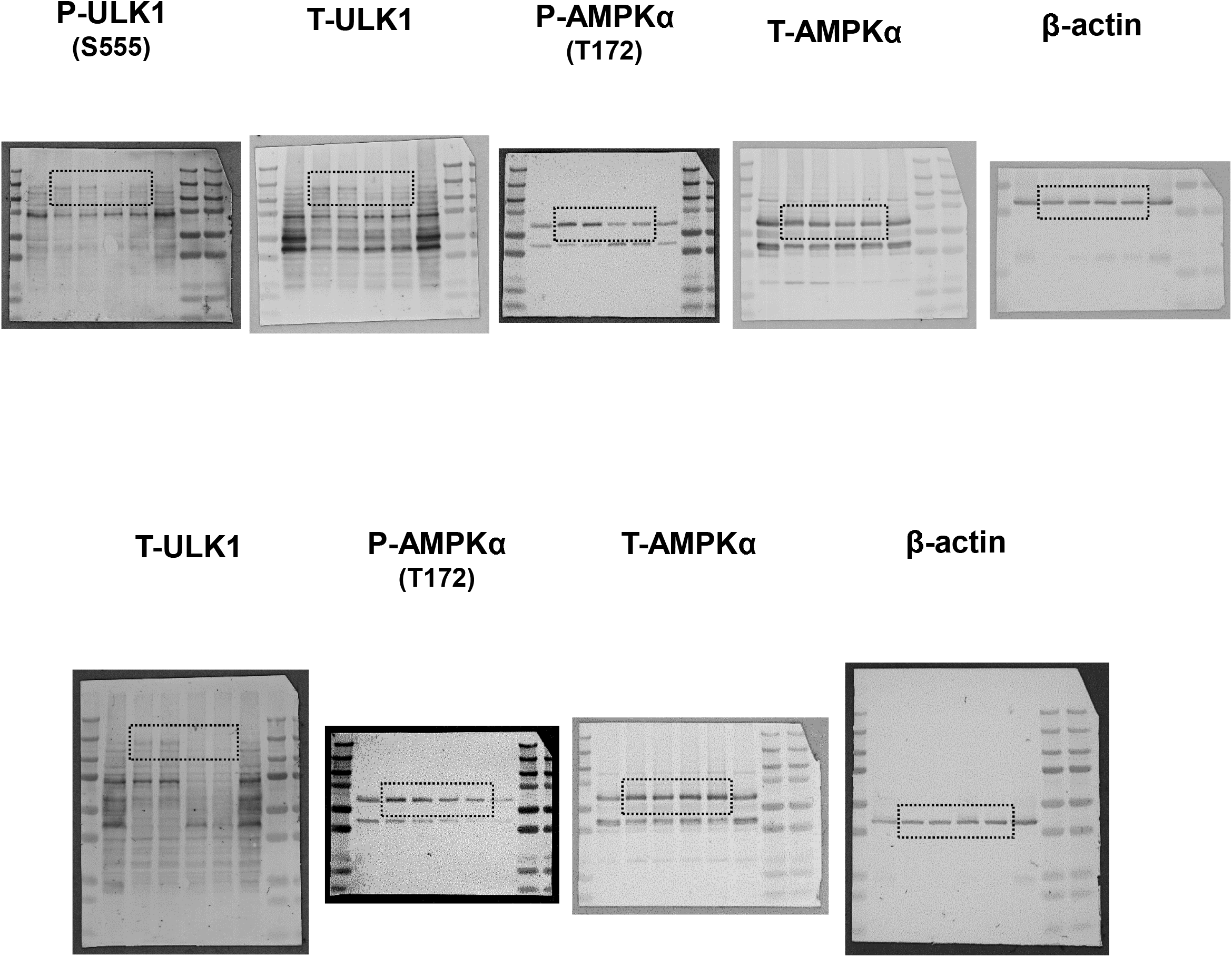

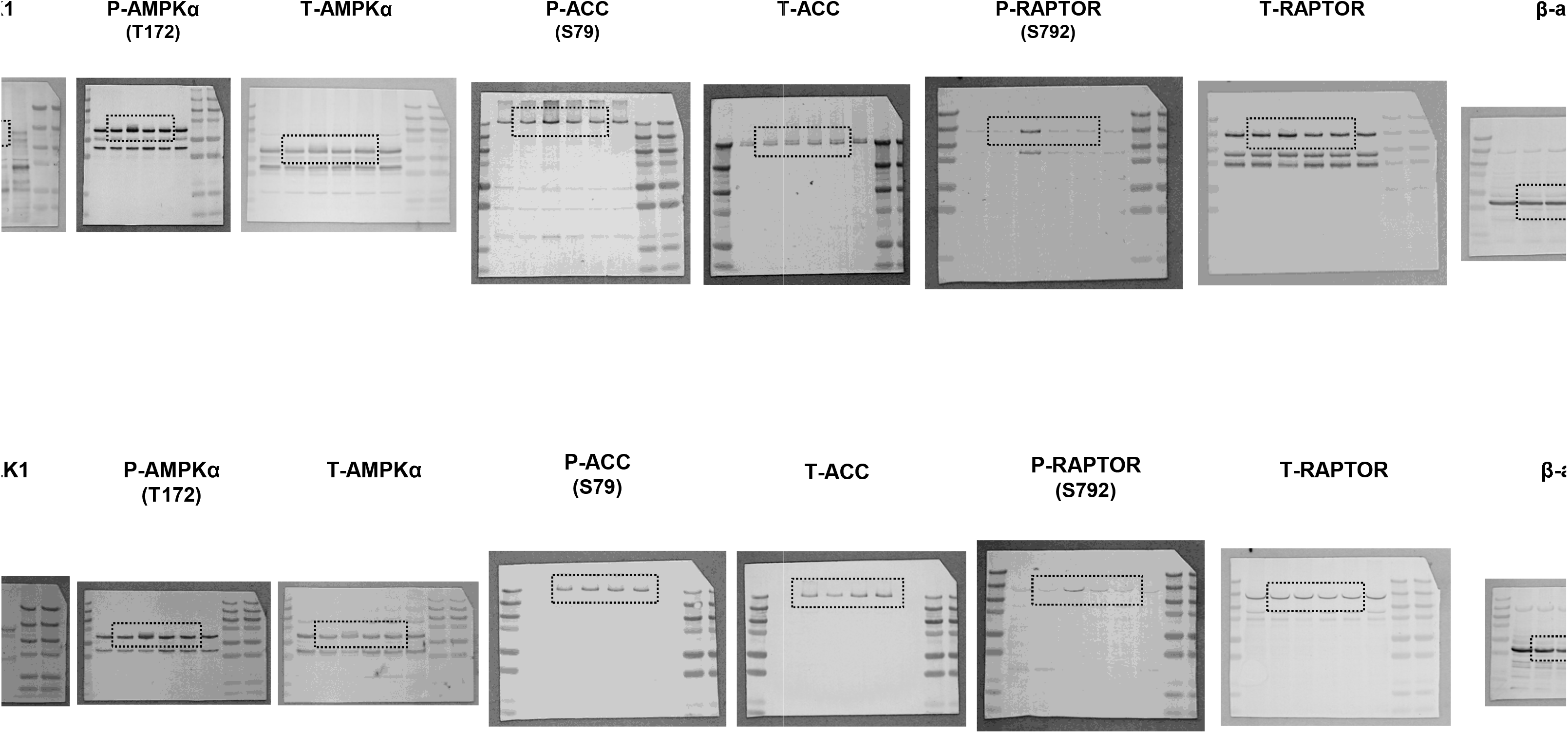

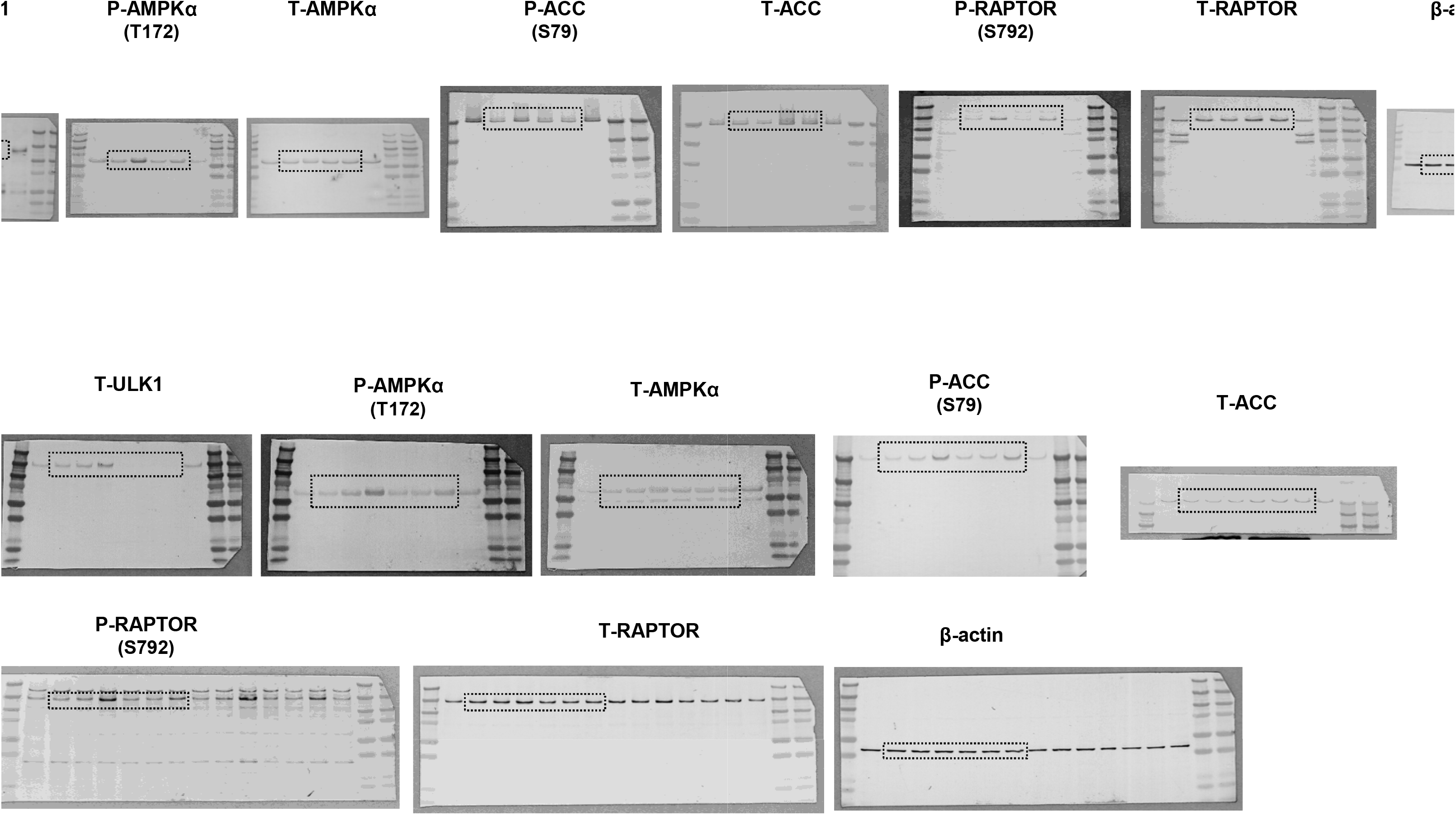

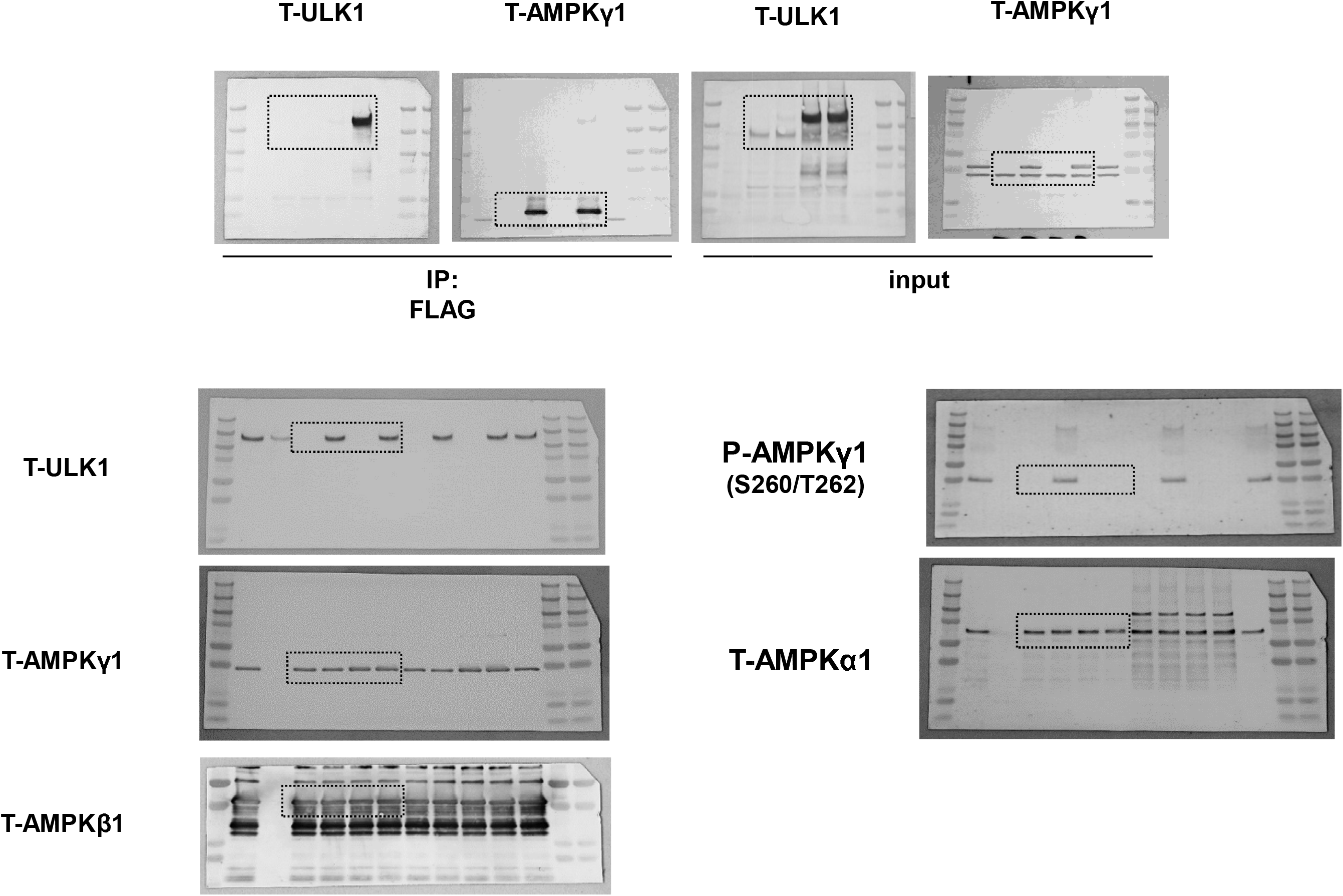

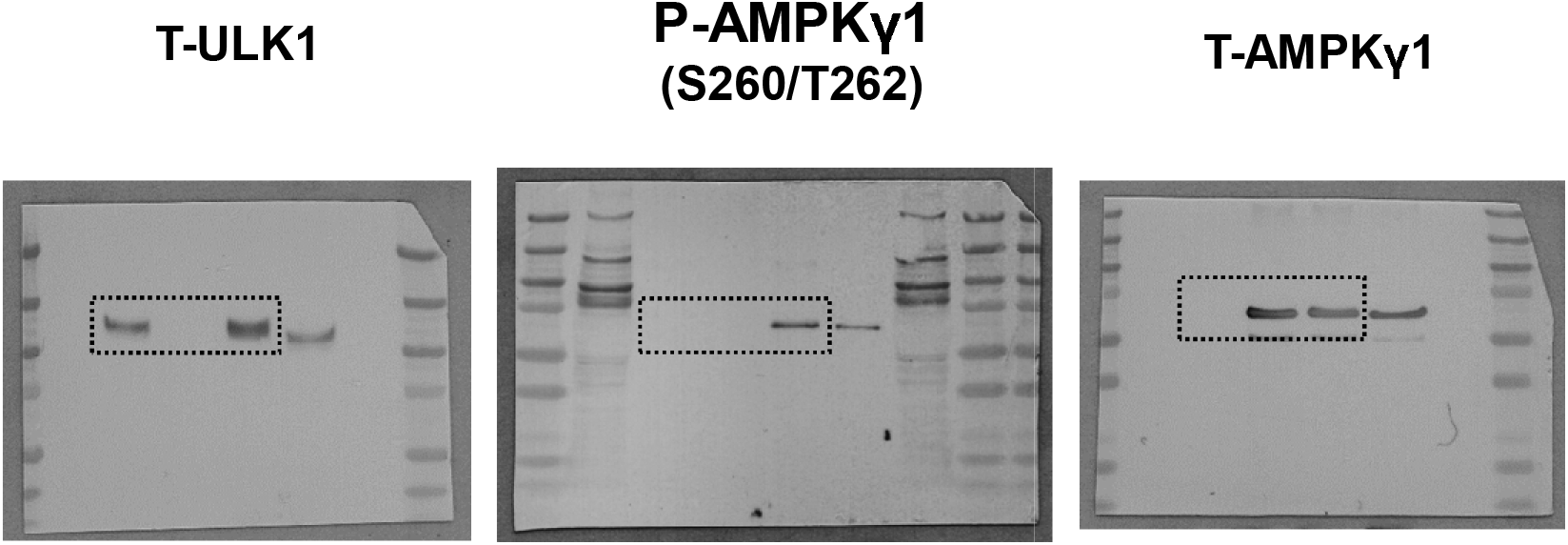

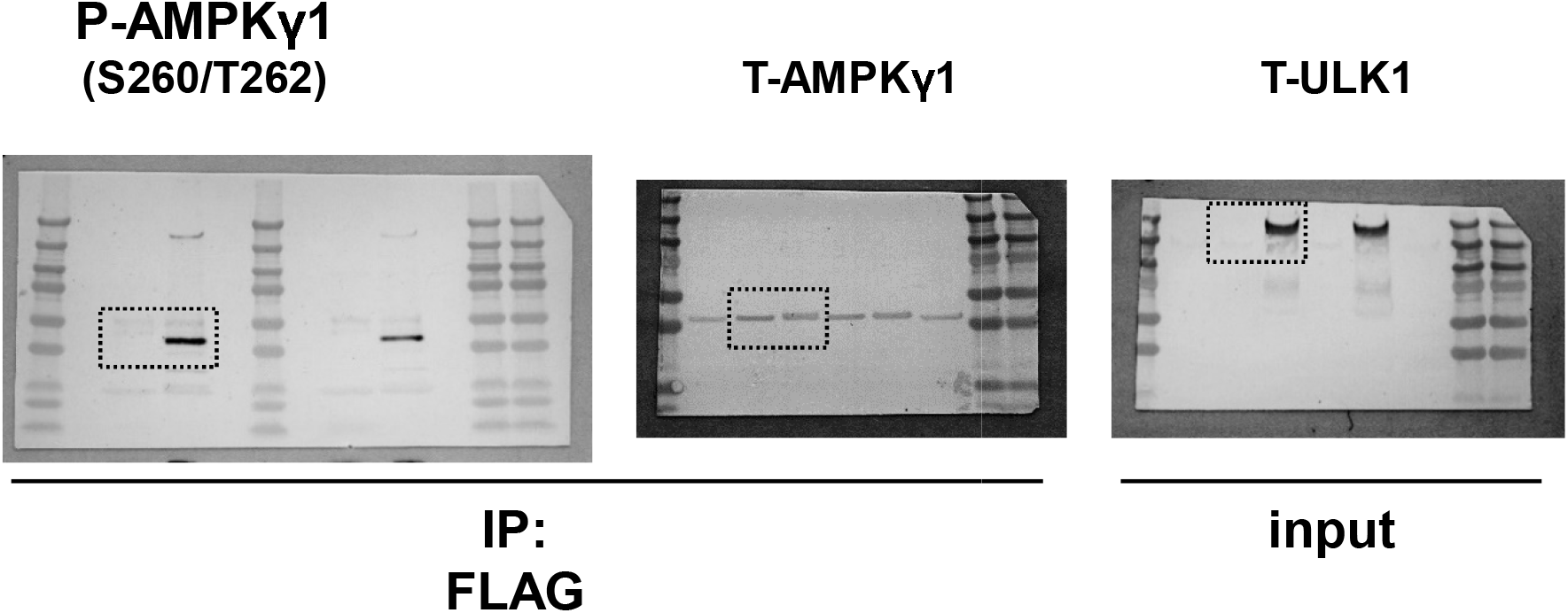

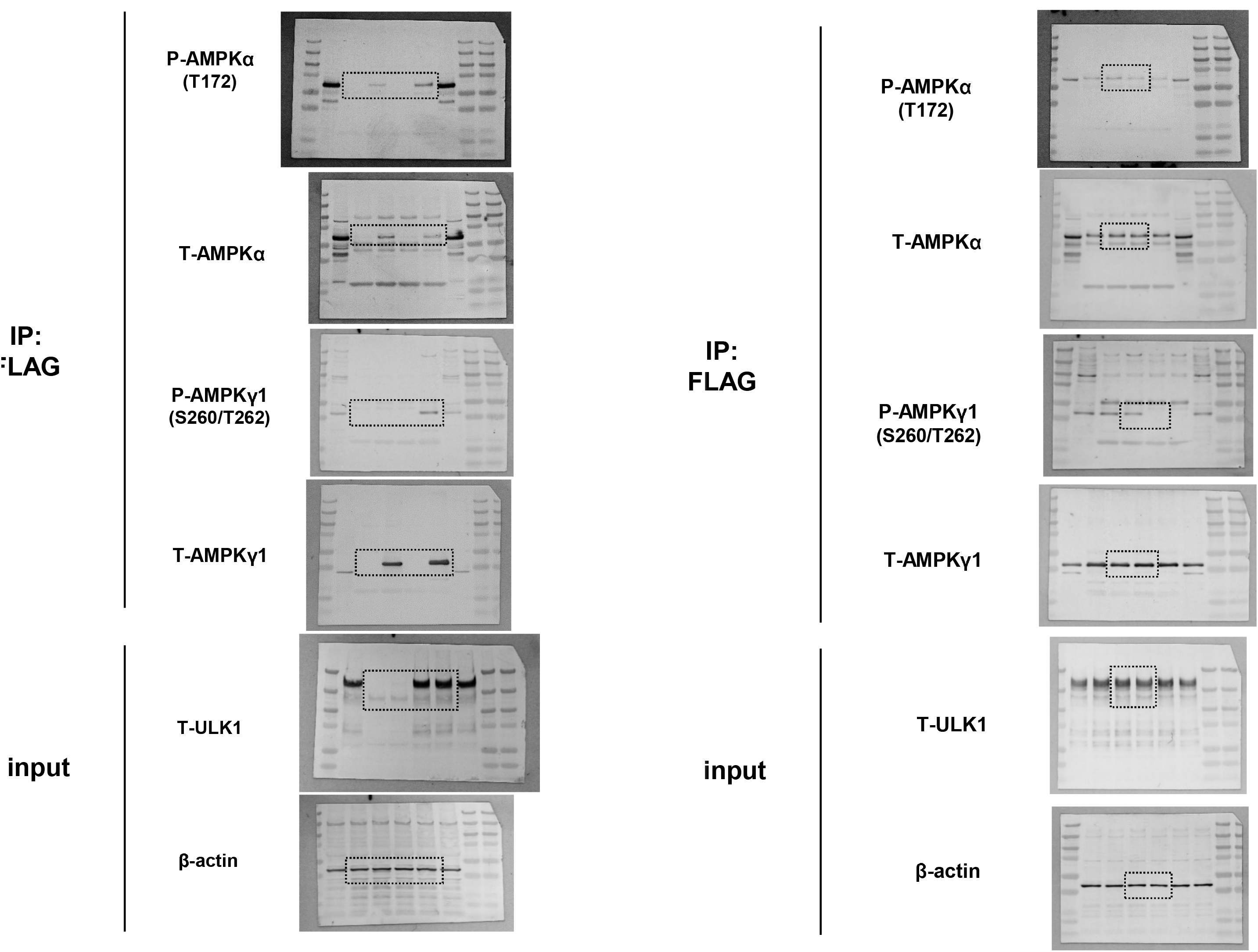

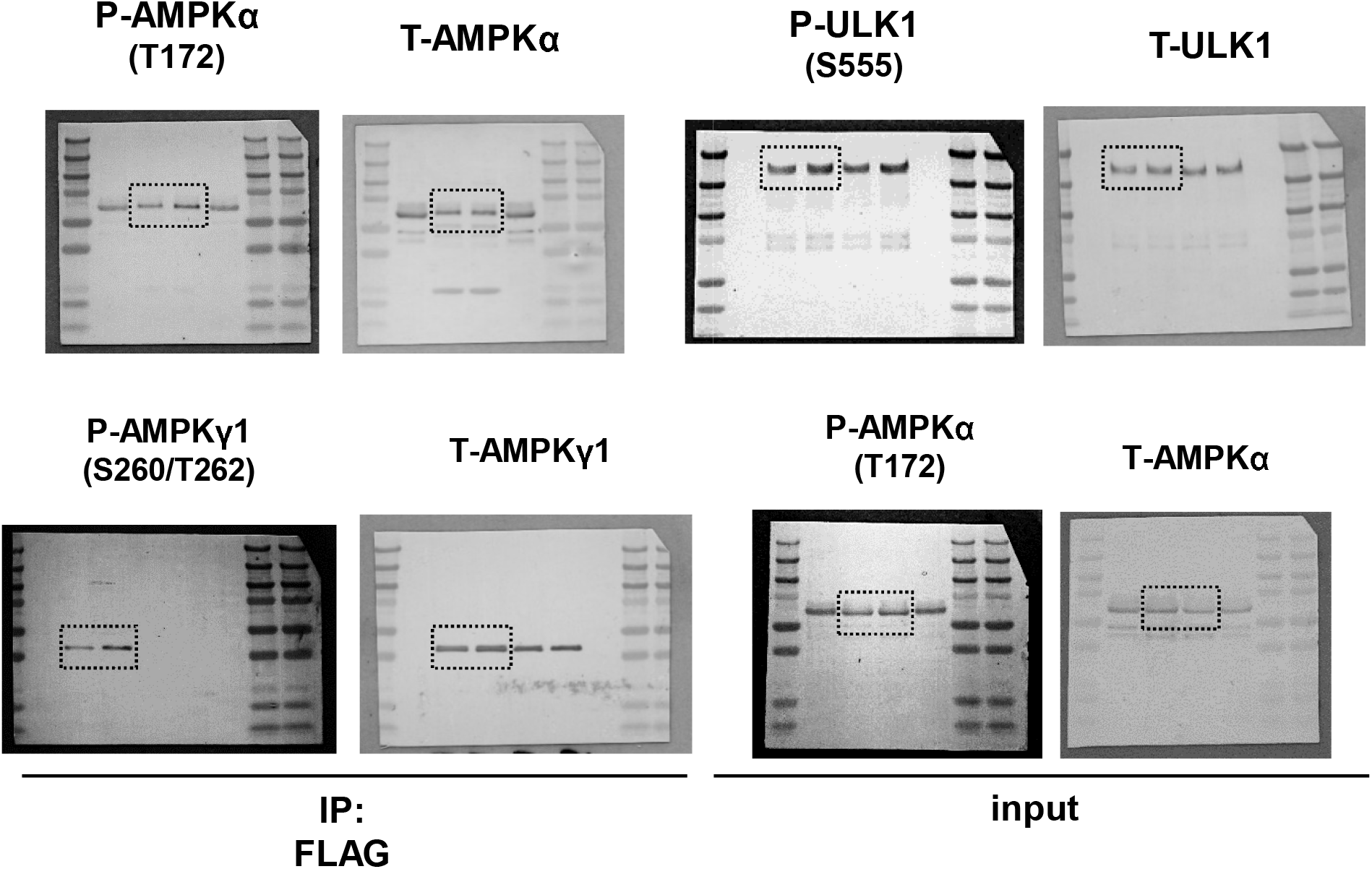

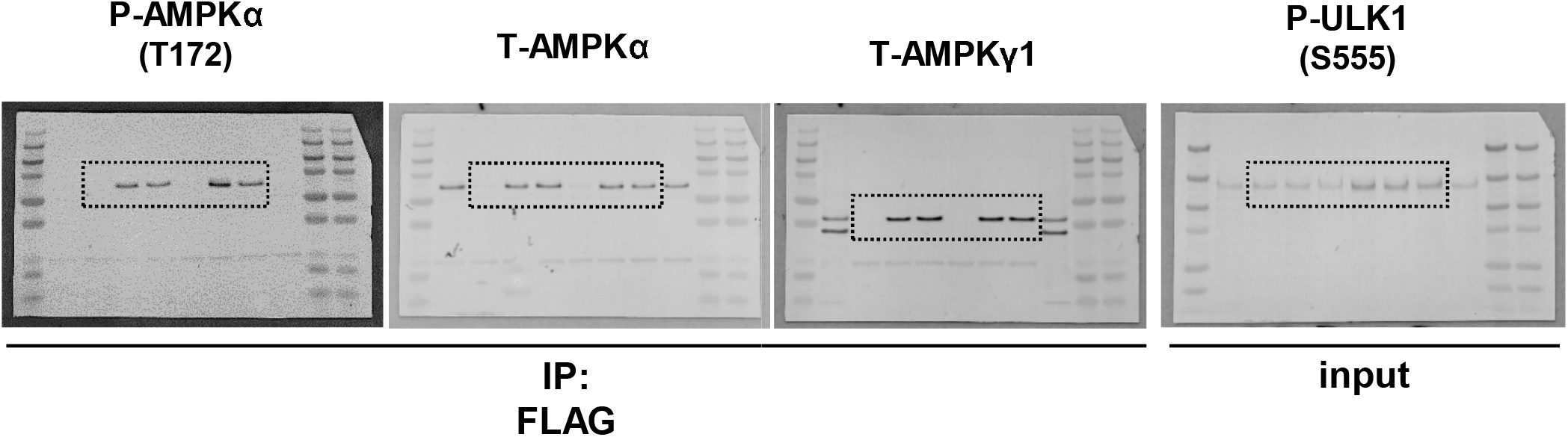

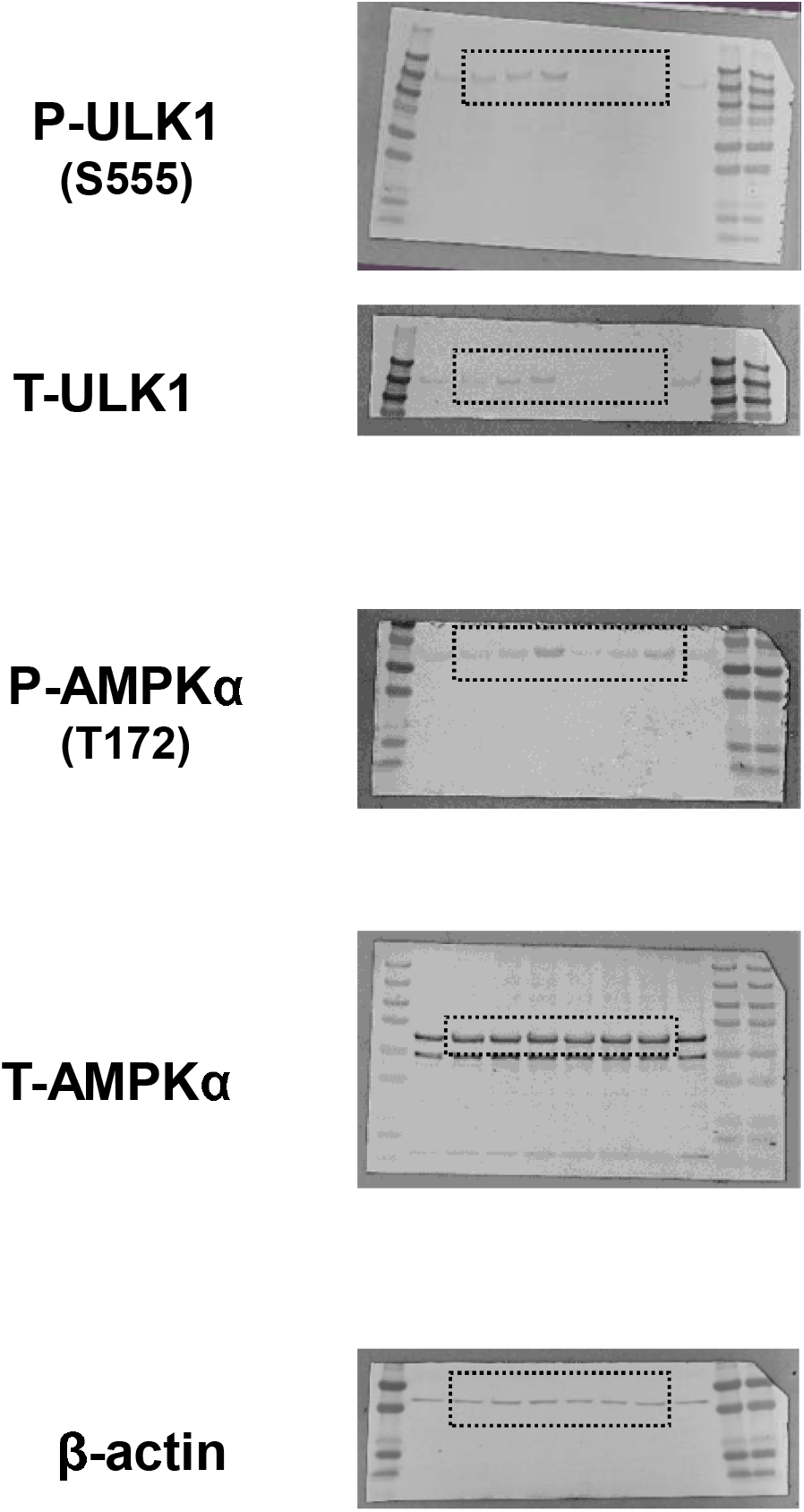

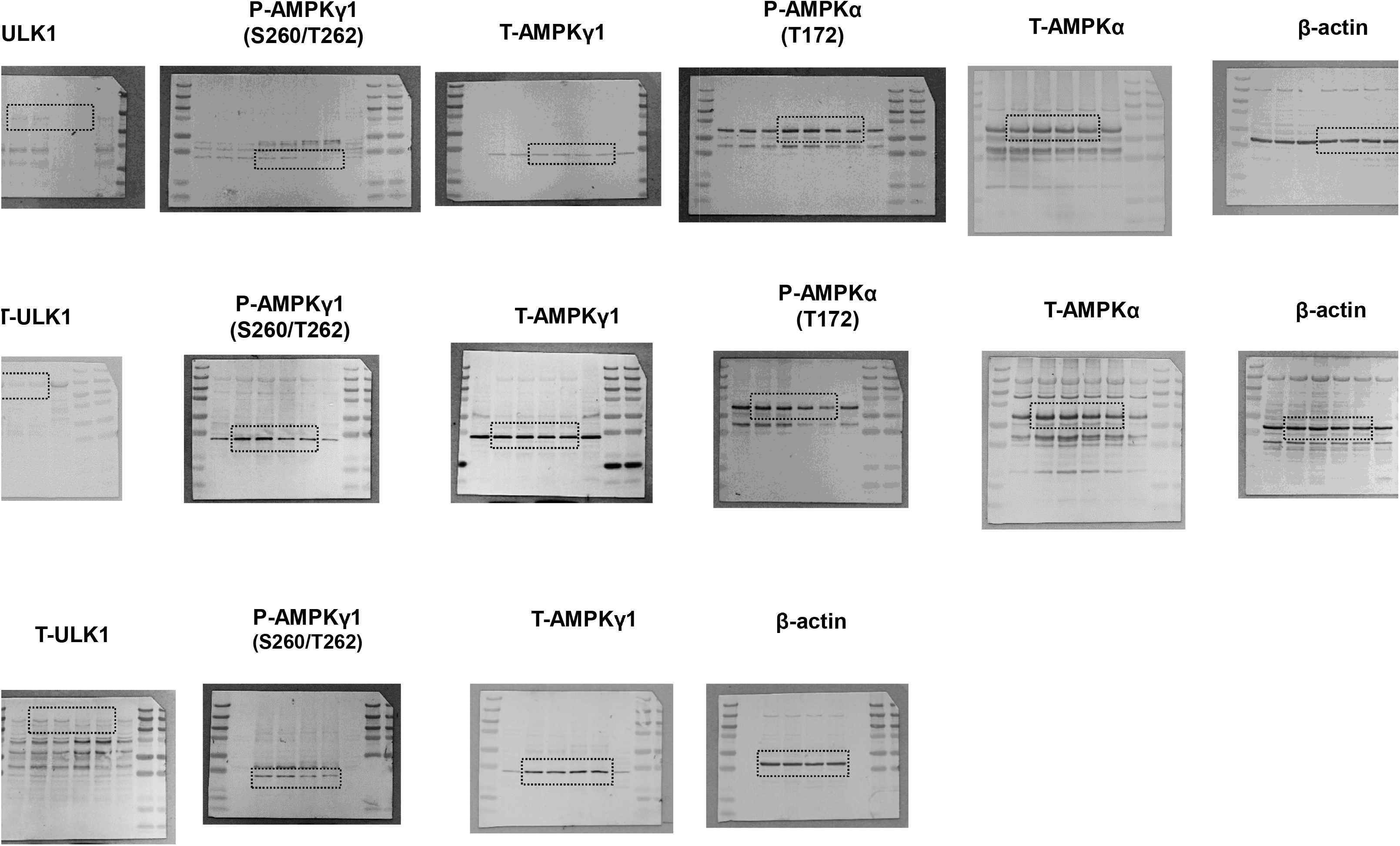

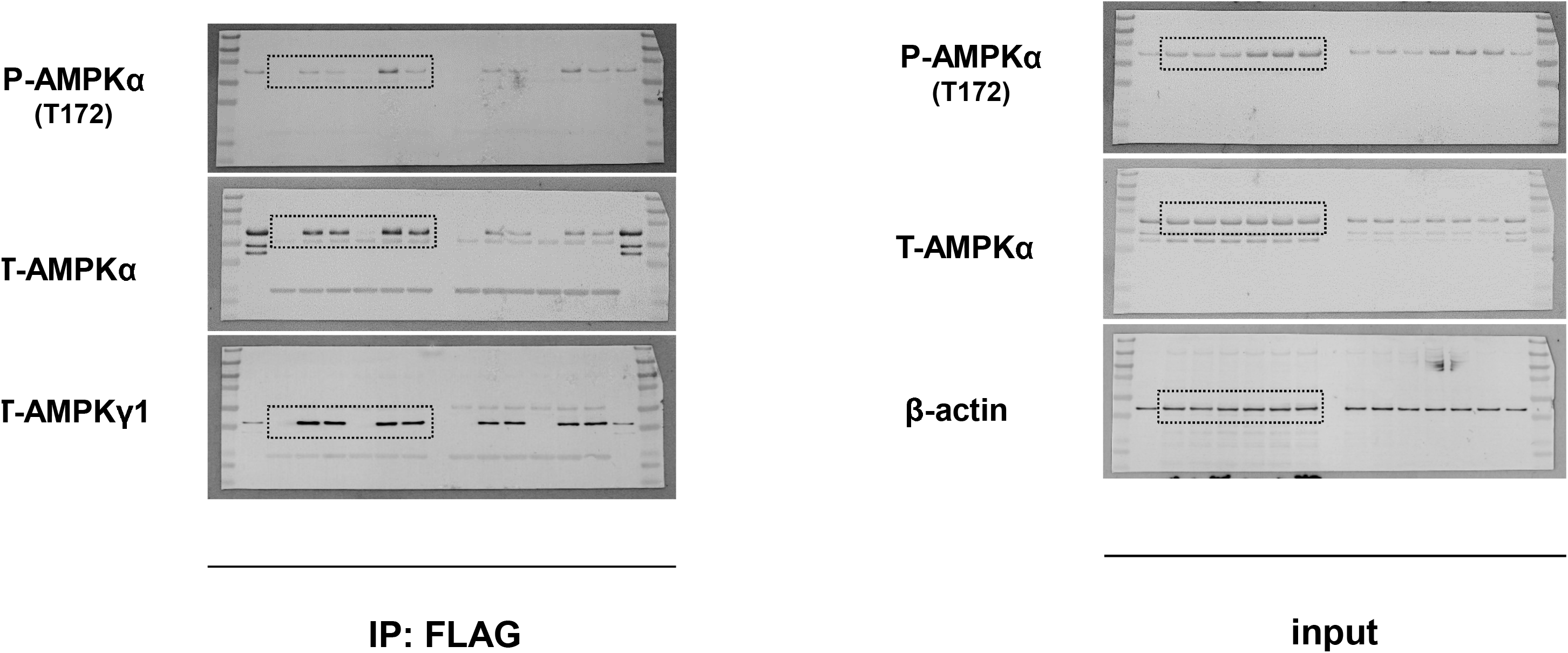

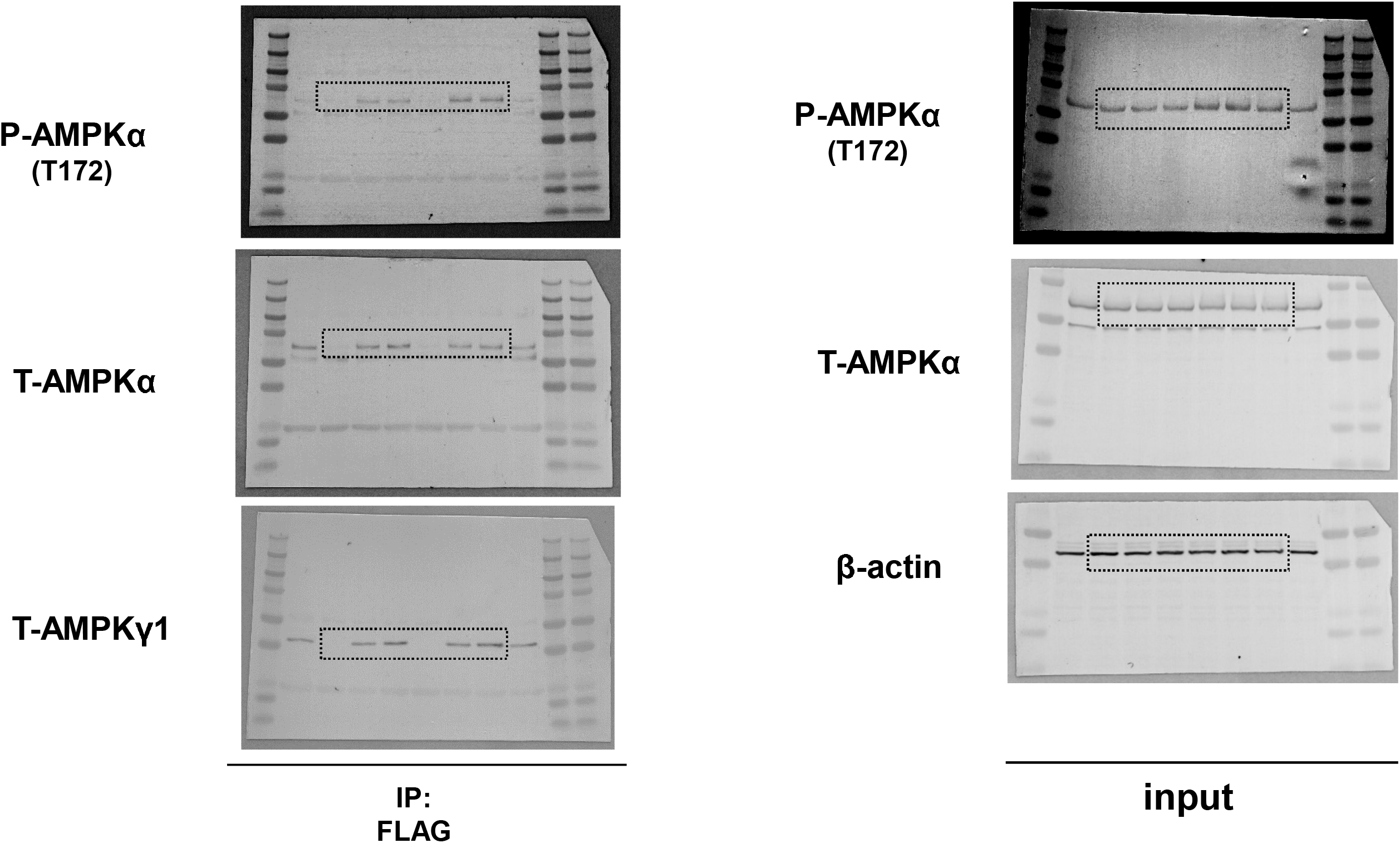

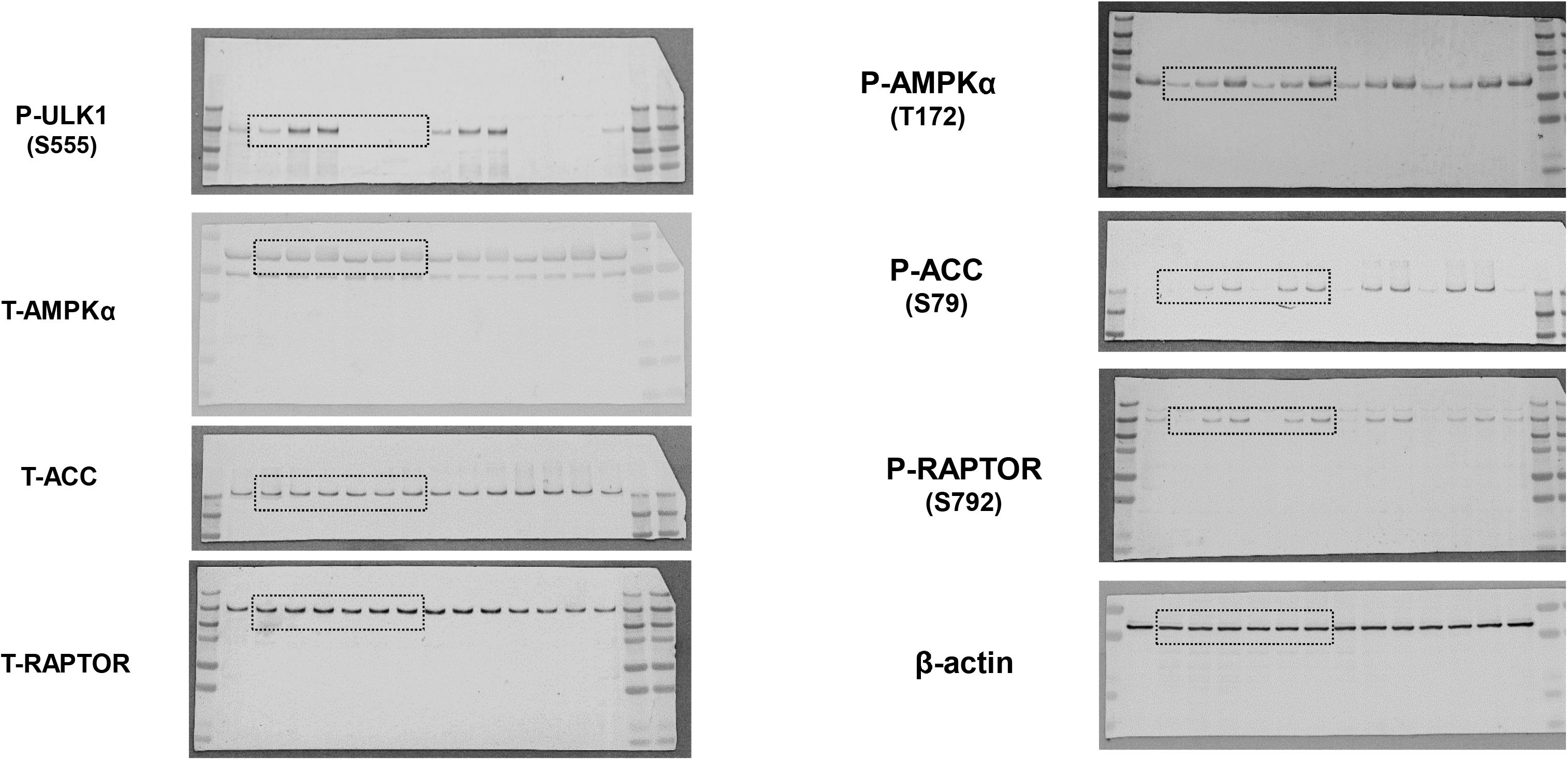

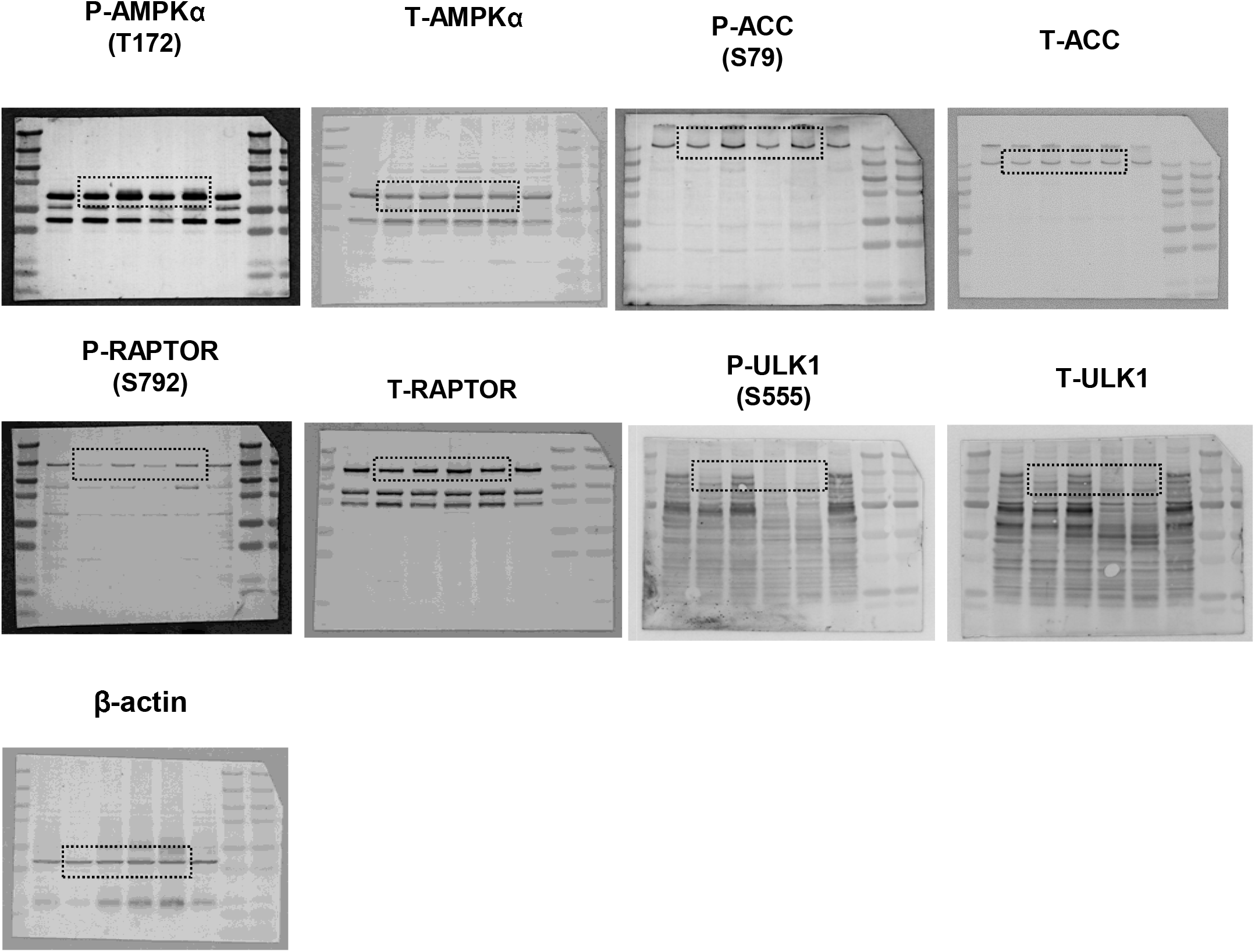

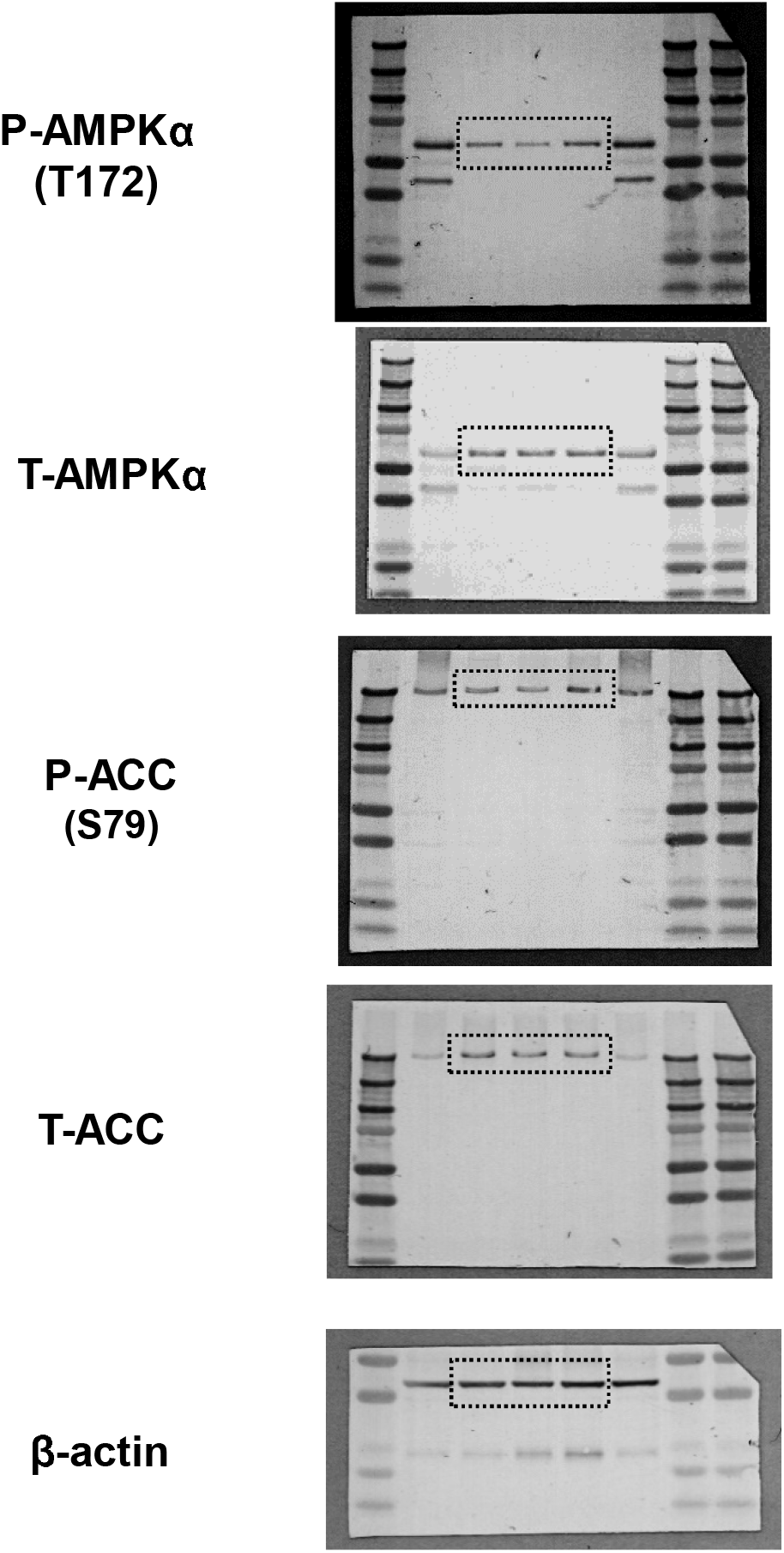

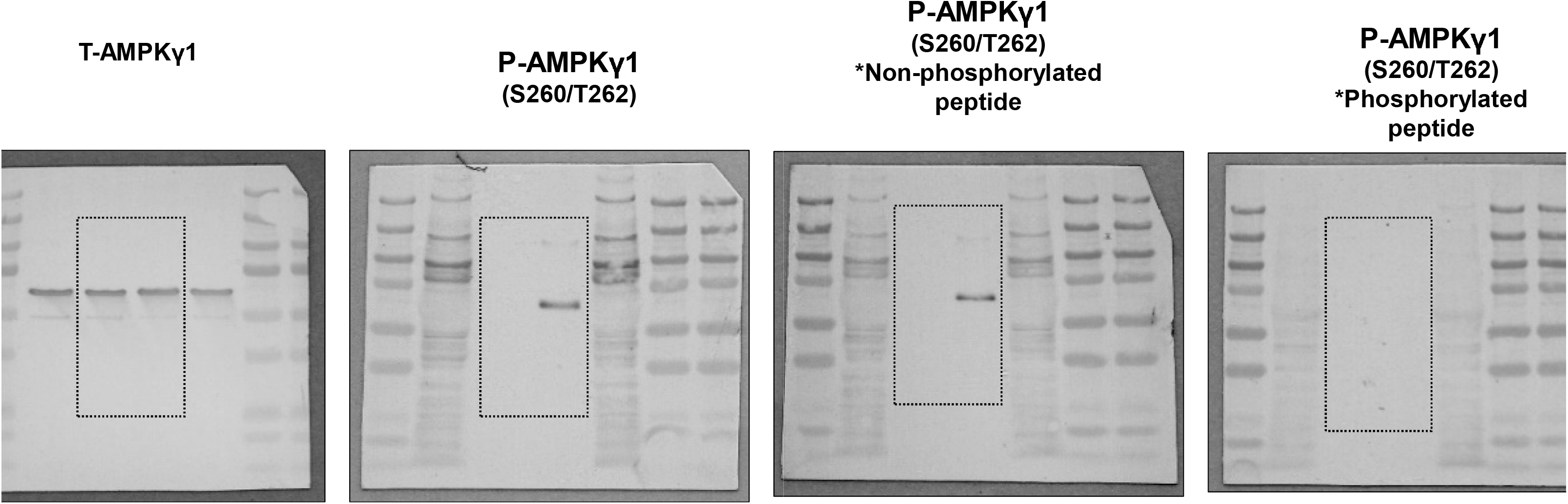

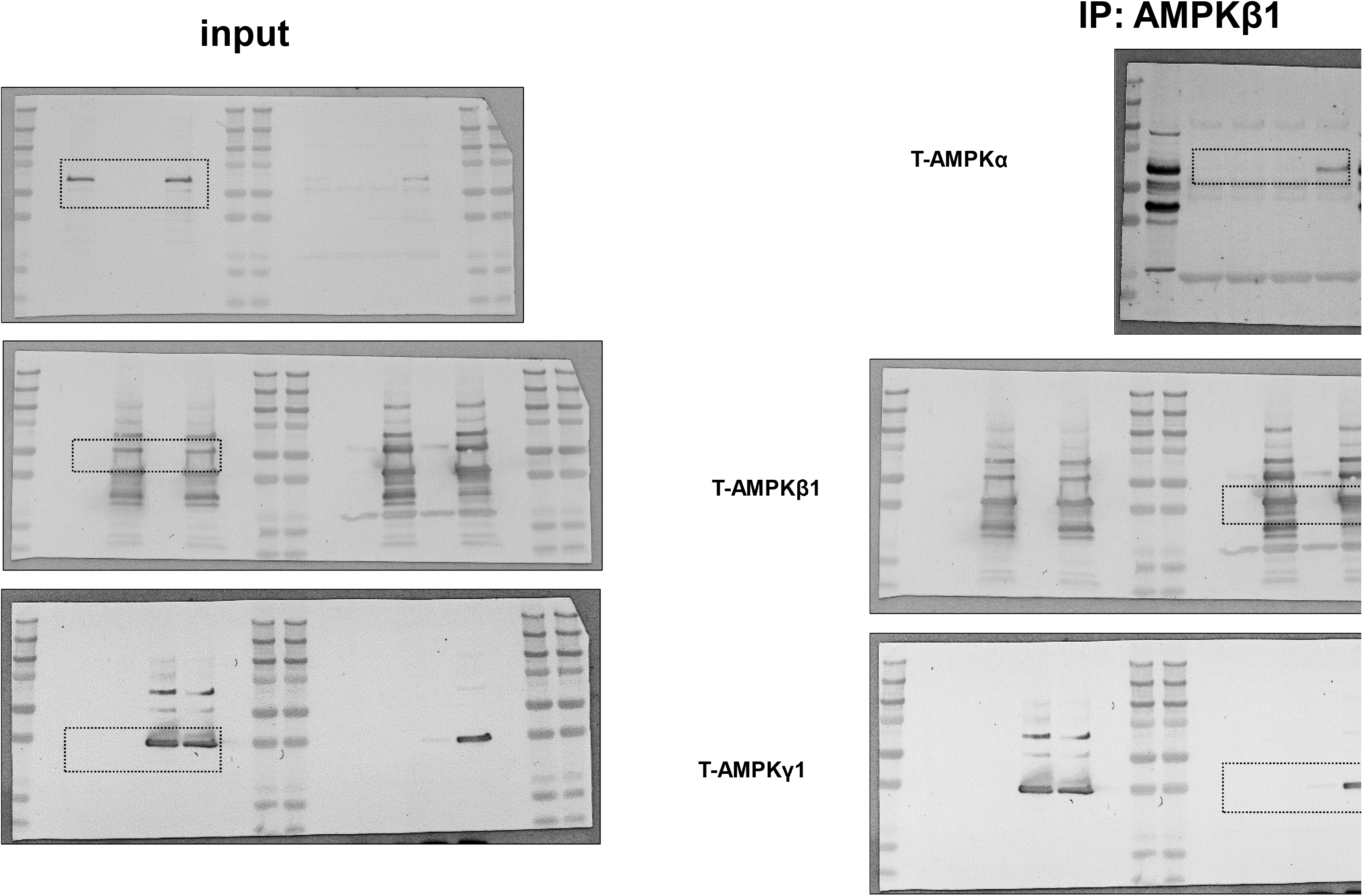

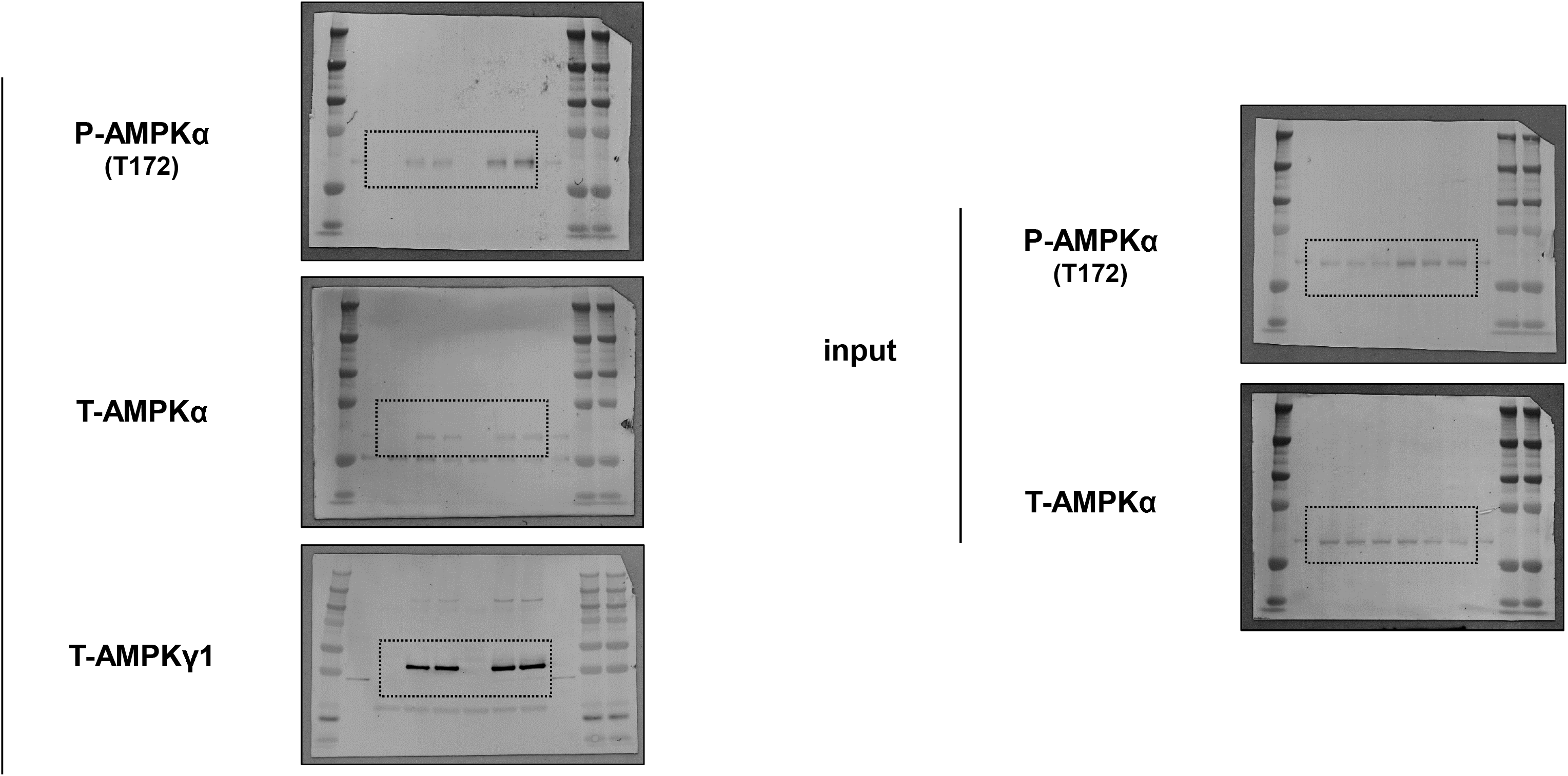

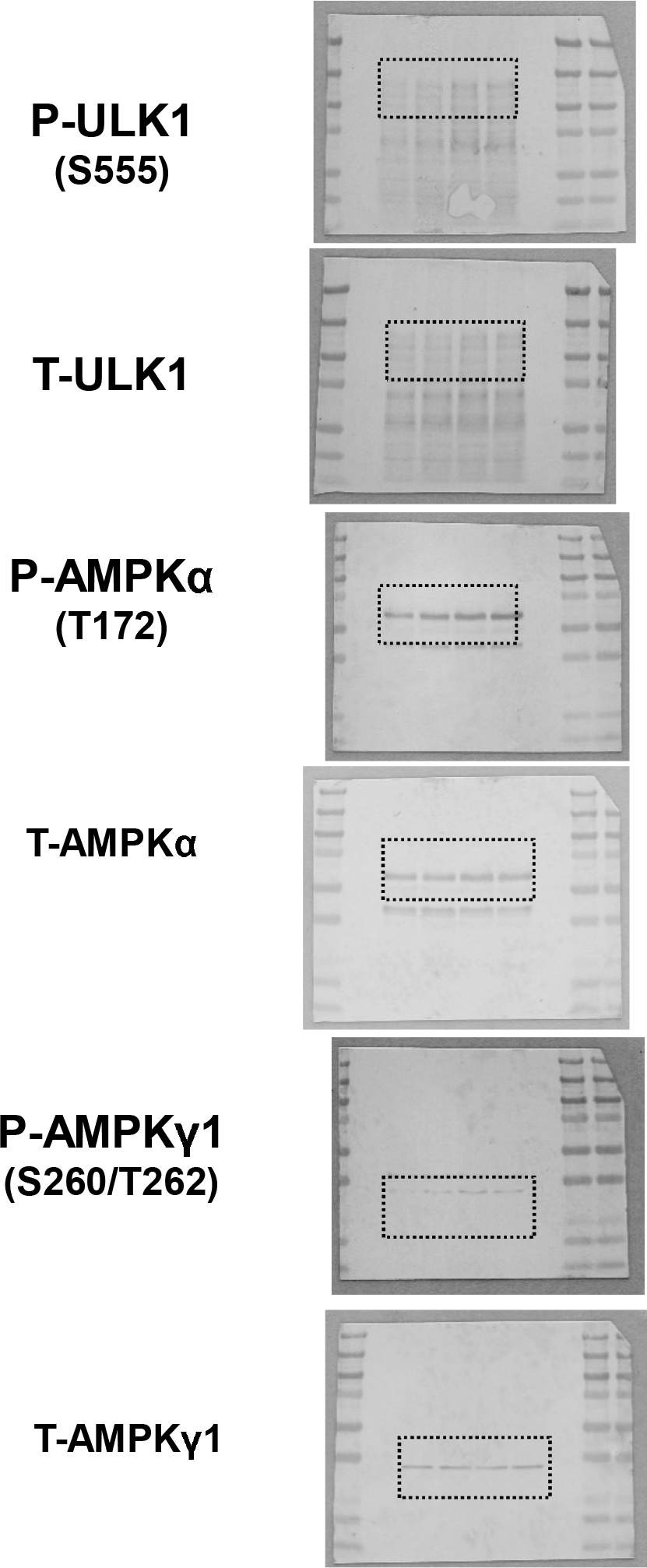

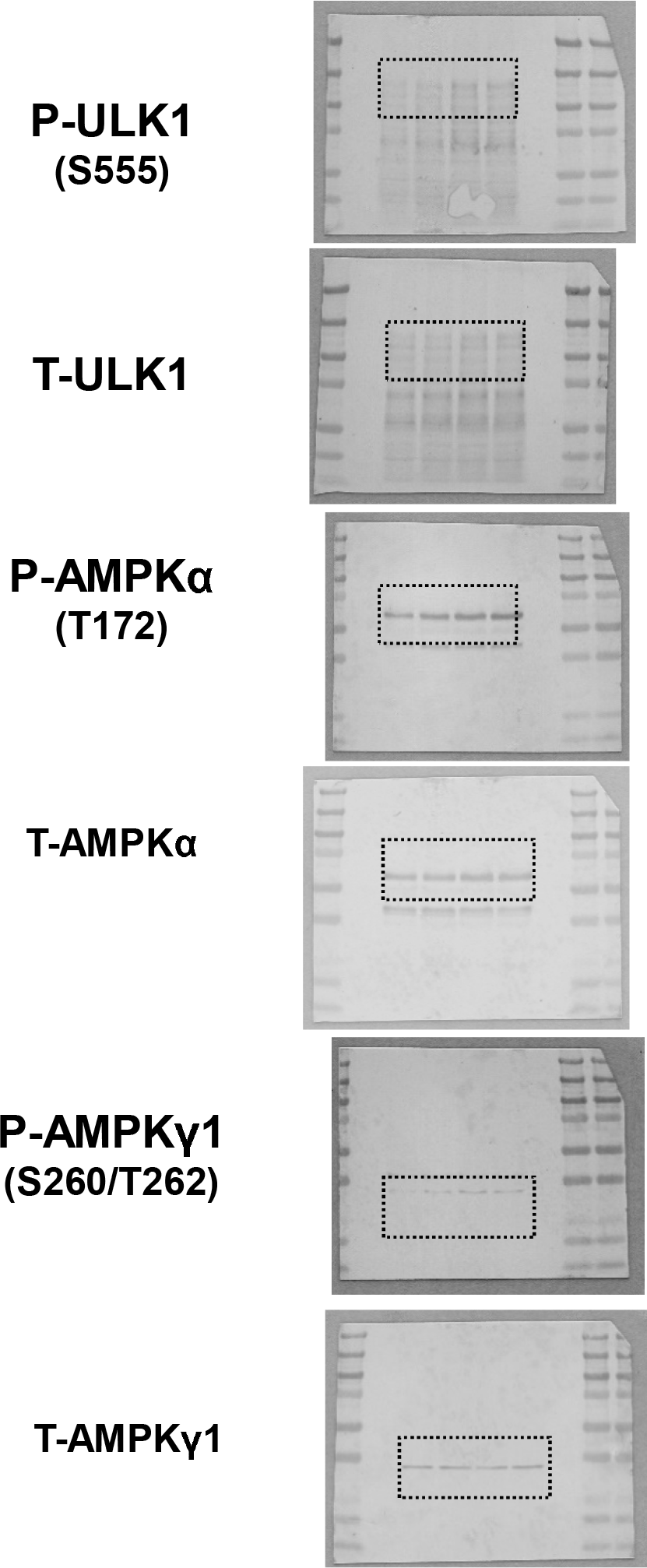

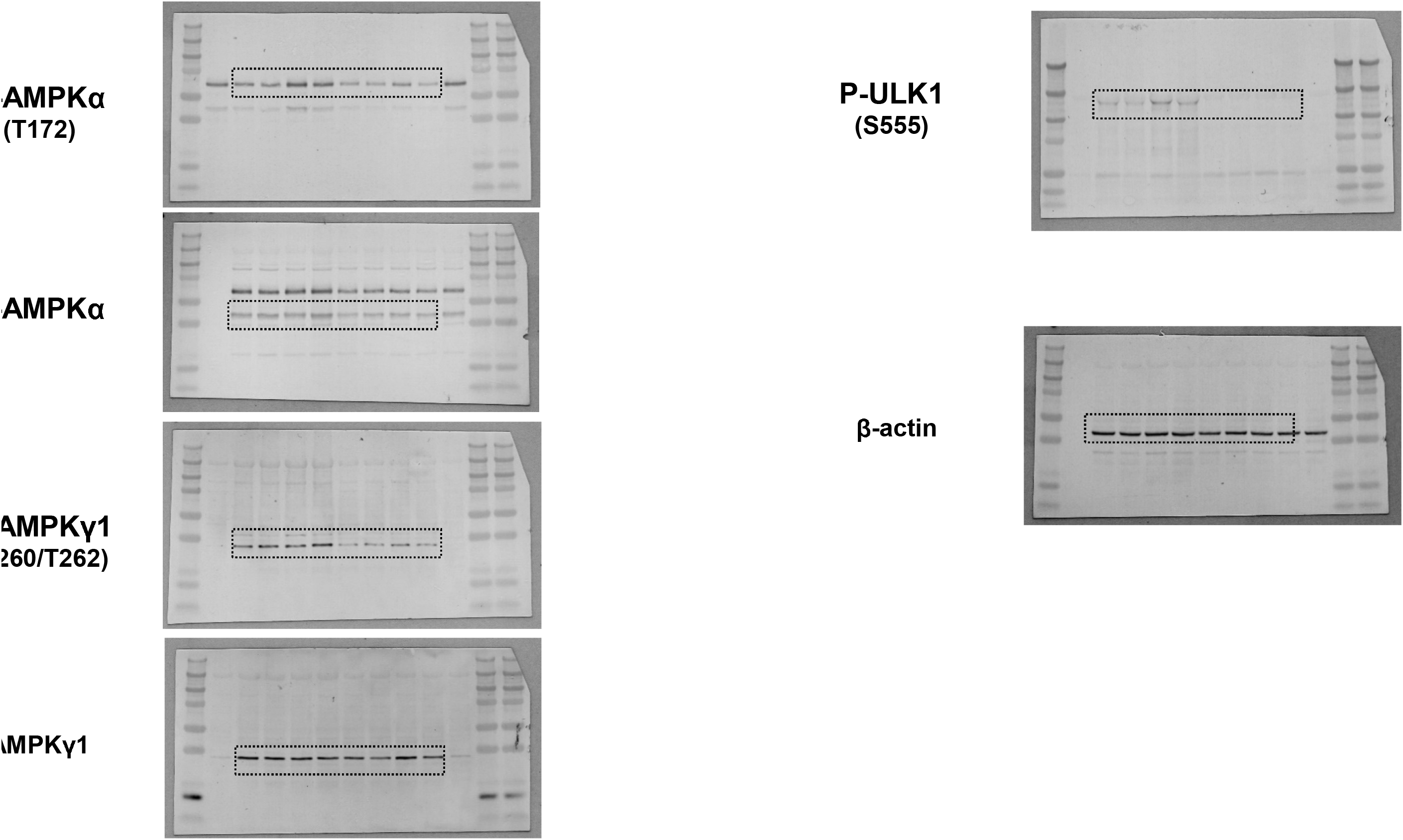

