## Supplementary Information for "ULK1-regulated AMP sensing by AMPK and its application for the treatment of chronic kidney disease"

**Supplementary method** : Statistical methods used are summarized in **source data file**.

**Supplemental figures**

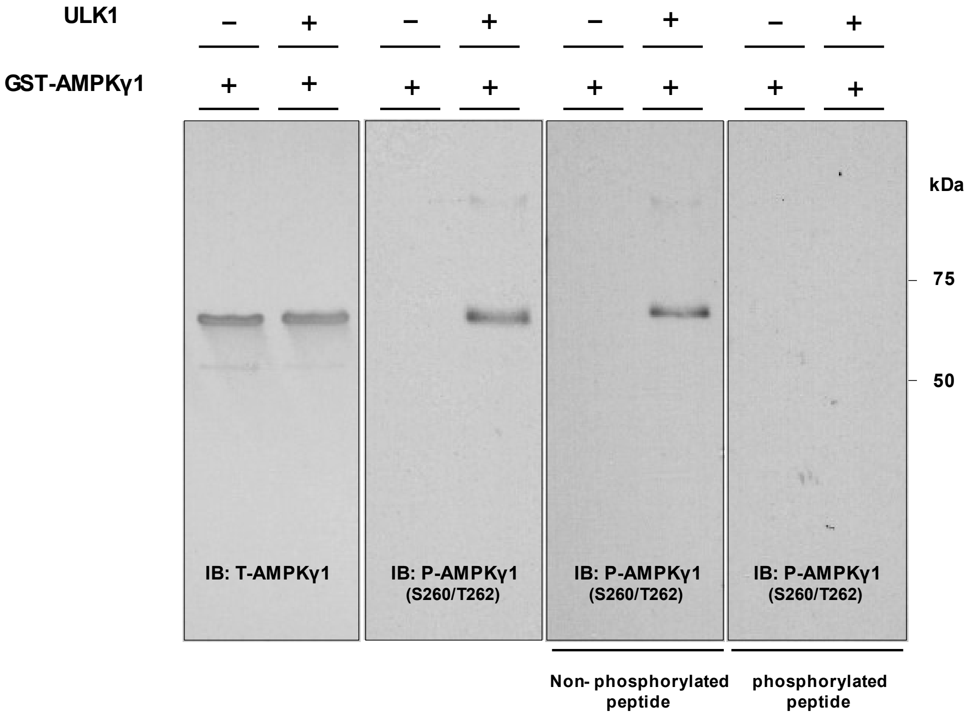


**Fig. S1. Evaluation of the generated anti-phospho-AMPKγ1$^{Ser260/Thr262}$ antibody**

The specificity of the antibody against phospho-AMPKγ1$^{Ser260/Thr262}$ was verified by an antigen absorption test. The band disappeared when the antibody was mixed with phosphorylated peptide, confirming the specificity of the antibody. AMPKγ1, AMP-activated protein kinase gamma-1.

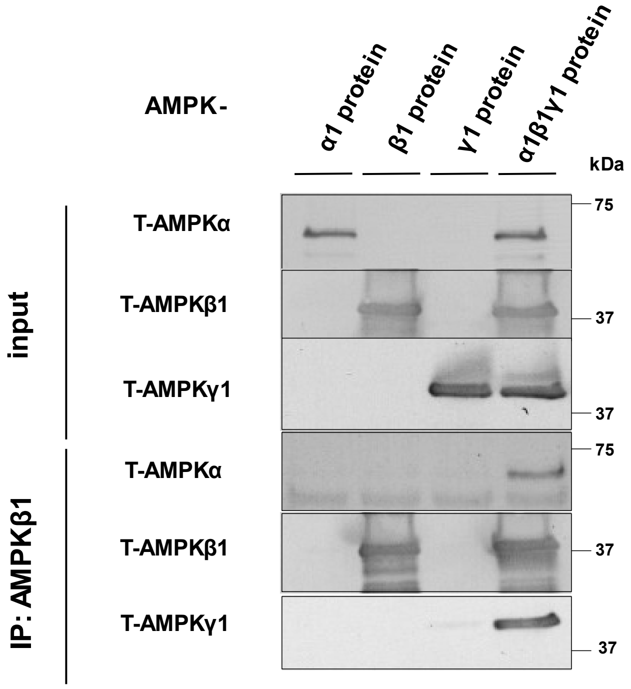


**Fig. S2. Evaluation of purified AMP-activated protein kinase (AMPK) trimeric protein**

Immunoblots of purified AMPKα1, AMPKβ1, AMPKγ1, and co-incubated AMPKα1β1γ1. Immunoprecipitation of AMPKα1β1γ1 with AMPKβ1 antibody confirmed the binding of AMPKα1 and AMPKγ1 to AMPKβ1 protein.

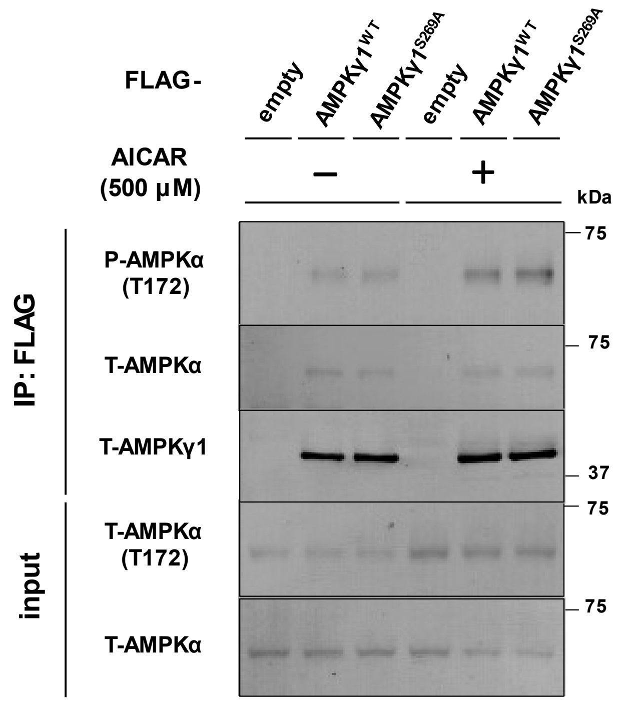


**Fig. S3. Evaluation of the sensitivity of P-AMPKα$^{Thr172}$ complexed with wild-type (WT) AMPKγ1 or S269A mutant to 5-aminoimidazole 4-carboxamide-1-beta-D-ribofuranoside (AICAR)**

Representative immunoblots evaluating the sensitivity of P-AMPKα$^{Thr172}$ complexed with wild-type (WT) AMPKγ1 or the S269A mutant to AICAR (500 μM), exhibiting no difference in the sensitivity to AICAR treatment between the WT and S269A mutant.

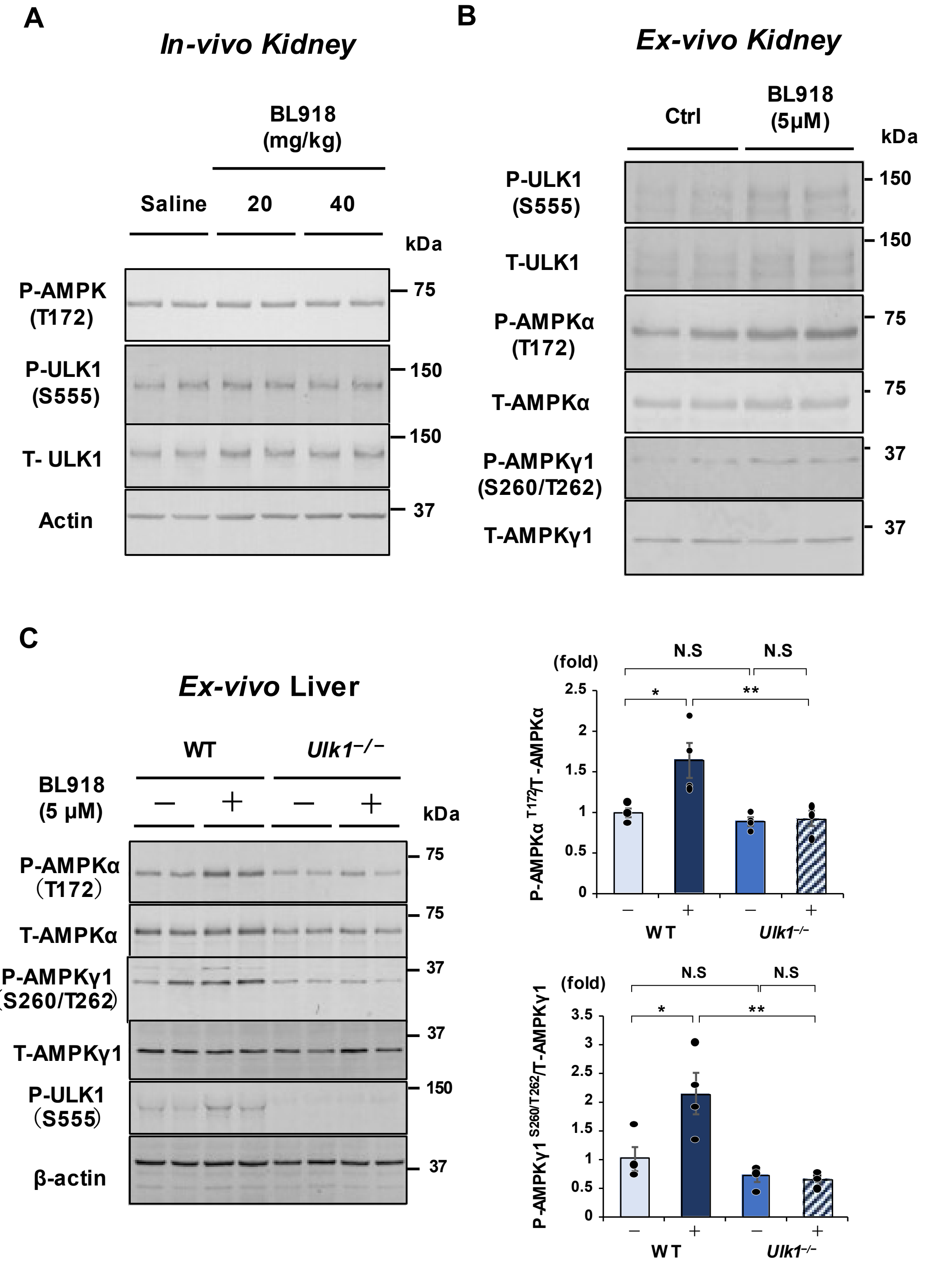


**Fig. S4. Evaluation of the effects of Unc-51-like kinase 1 (ULK1)-selective activator (BL918) on AMP-activated protein kinase (AMPK) in the kidneys and liver**

**A**. Representative immunoblots of P-AMPKα$^{Thr172}$ and P-ULK1 $^{Ser555}$ in the kidneys treated with either saline or BL918 (20 mg/kg or 40 mg/kg) showing no significant difference in P-AMPKα$^{Thr172}$ phosphorylation between BL918 treatment and saline control.

**B.** Representative immunoblots of P-AMPKα$^{Thr172}$ and P-ULK1 $^{Ser555}$ in *ex-vivo* kidney tissues treated with BL918 (5 μM) relative to kidney tissues treated without BL-918.

**C.** Representative immunoblots (Left) and densitometric analysis (Right) of P-AMPKα$^{Thr172}$ and P-AMPKγ1$^{Ser260/Thr262}$ expression in *ex-vivo* liver tissues of wild-type (WT) or *Ulk1*$^{-/-}$ mice treated with BL918 (5 μM) relative to liver tissues treated without BL-918 (n = 4).

*P* values (**P* < 0.05, ***P* < 0.01) were calculated using the two-tailed unpaired *t*-test or one-way analysis of variance with Tukey's honestly significant difference test. Data are presented as the mean ± standard error of the mean. For detailed data, statistical analysis, and exact P values, refer to the **Source Data file**.

| | Residues | Sequence |
| --- | --- | --- |
| Mouse AMPK γ1 | 260, 262 | Y N N L D V **S** V **T** K A L Q H R S |
| Mouse AMPK γ2 | 490, 492 | Y N N L D I **T** V **T** Q A L Q H R S |
| Human AMPK γ1 | 270, 272 | Y N N L D V **S** V **T** K A L Q H R S |
| Human AMPK γ2 | 265, 267 | Y N N L D I **T** V **T** Q A L Q H R S |

**Fig. S5. Conserved sequence alignment between AMPKγ1 and AMPKγ2**

### Supplementary Tables

| Complex, Treatment | Fluorescence intensity | | | | | | |
|---|---|---|---|---|---|---|---|
| | **Batch 1** | **Batch 2** | **Batch 3** | **Batch 4** | **Batch 5** | **Average** | **SD** |
| **$\alpha 1\beta 1\gamma 1^{WT}$** | 1.000 | 1.000 | 1.000 | 1.000 | 1.000 | **1.000** | 0.000 |
| **$\alpha 1\beta 1\gamma 1^{WT}$ + ULK1** | 1.074 | 1.141 | 1.239 | 1.466 | 2.080 | **1.400** | 0.408 |
| **$\alpha 1\beta 1\gamma 1^{S260A/T262A}$** | 0.565 | 0.695 | 0.443 | 0.480 | 0.674 | **0.571** | 0.113 |
| **$\alpha 1\beta 1\gamma 1^{S260A/T262A}$ + ULK1** | 0.484 | 0.434 | 0.635 | 0.635 | 0.729 | **0.584** | 0.121 |

**Supplementary Table 1**

**Measured fluorescence intensity of Mant-AMP bound to trimeric AMPK$\alpha 1^{WT}\beta 1^{WT}\gamma 1^{WT}$ (AMPK$\alpha 1\beta 1\gamma 1^{WT}$) or AMPK$\alpha 1^{WT}\beta 1^{WT}\gamma 1^{S260A/T262A}$ mutant (AMPK$\alpha 1\beta 1\gamma 1^{S260A/T262A}$) protein complex co-incubated with or without ULK1.**

All intensity values are normalized by the intensity detected in AMPK$\alpha 1\beta 1\gamma 1^{WT}$ without ULK1 co-incubation.

| Gene name | Forward | Reverse |
|---|---|---|
| *Gapdh* | TGTGTCCGTCGTGGATCTGA | TTGCTGTTGAAGTCGCAGGAG |
| *Col1a* | GCTCCTCTTAGGGGCCACT | CCACGTCTCACCATTGGGG |
| *Col3a* | AACCTGGTTTCTTCTCACCCTTC | ACTCATAGGACTGACCAAGGTGG |
| *Fn1* | ACAAGGTTCGGGAAGAGGTT | CCGTGTAAGGGTCAAAGCAT |
| *Acta2* | CTGACAGAGGCACCACTGAA | AGAGGCATAGAGGGACAGCA |

**Supplementary Table 2**

**Sequences of oligonucleotide primers used for quantitative polymerase chain reaction (qRT-PCR)**

*Col1a,* collagen type 1 alpha; *Col3a*; collagen type 3 alpha; *Fn1*, fibronectin 1; *Acta2*; actin alpha-2
